# TBK1 is ubiquitinated by TRIM5α to assemble mitophagy machinery

**DOI:** 10.1101/2023.10.19.563195

**Authors:** Bhaskar Saha, Hallvard Olsvik, Geneva L Williams, Seeun Oh, Gry Evjen, Eva Sjøttem, Michael A Mandell

## Abstract

Ubiquitination of mitochondrial proteins provides a basis for the downstream recruitment of mitophagy machinery, yet whether ubiquitination of the machinery itself contributes to mitophagy is unknown. Here, we show that K63-linked polyubiquitination of the key mitophagy regulator TBK1 is essential for its mitophagy functions. This modification is catalyzed by the ubiquitin ligase TRIM5α. Mitochondrial damage triggers TRIM5α’s auto-ubiquitination and its interaction with ubiquitin-binding autophagy adaptors including NDP52, optineurin, and NBR1. Autophagy adaptors, along with TRIM27, enable TRIM5α to engage with TBK1. TRIM5α with intact ubiquitination function is required for the proper accumulation of active TBK1 on damaged mitochondria in Parkin-dependent and Parkin-independent mitophagy pathways. Additionally, we show that TRIM5α can directly recruit autophagy initiation machinery to damaged mitochondria. Our data support a model in which TRIM5α provides a self-amplifying, mitochondria-localized, ubiquitin-based, assembly platform for TBK1 and mitophagy adaptors that is ultimately required to recruit the core autophagy machinery.

## INTRODUCTION

Removal of unwanted or dysfunctional mitochondria is essential for cellular health. This is accomplished through mitophagy, an autophagy-based system for the selective degradation of mitochondria in lysosomes. Mitophagy is activated in response to mitochondrial damage or for the elimination of mitochondria as part of cellular development (e.g. reticulocyte maturation) (Onishi et al., 2021; Pickles et al., 2018). Alterations in mitophagy have been linked with cardiovascular, metabolic, or neurodegenerative diseases and cancer (Evans and Holzbaur, 2019; Gong et al., 2015; Pickrell et al., 2015). As such, there has been intensive interest in uncovering the molecular mechanisms underlying how cells selectively identify mitochondria for mitophagy-based elimination (Uoselis et al., 2023).

Several non-redundant mitophagy pathways have evolved. The molecular details of how mitochondria are identified by these pathways differ; nevertheless, they all involve the sequential and self-amplifying localization of a conserved set of autophagy proteins to mitochondria (Bunker et al., 2023; Vargas et al., 2019). Elegant studies have shown that the recruitment of the ULK1/FIP200 complex to mitochondria is necessary and sufficient for mitophagy induction (Bunker et al., 2023; Hung et al., 2021; Lazarou et al., 2015; Vargas et al., 2019). This complex can stimulate the mitochondrial localization and activation of downstream factors required for the initiation and execution of autophagosome formation. The optimal recruitment of the ULK1/FIP200 complex to damaged mitochondria is dependent on the accumulation or presentation of proteins on the mitochondria’s cytoplasmic face that mark it for recognition by the autophagy machinery (Abudu et al., 2021; Heo et al., 2015; Kane et al., 2014; Princely Abudu et al., 2019; Wei et al., 2017). Ubiquitination of proteins on the outer mitochondrial membrane is one such marker (Harper et al., 2018; Zachari et al., 2019). This ubiquitin can serve as a ligand for ubiquitin-binding adaptor proteins that interact with the ULK1/FIP200 complex and other autophagy proteins (e.g. LC3s and GABARAPs) and scaffold them onto the surface of the ubiquitinated mitochondria (Heo et al., 2015; Lazarou et al., 2015; Ravenhill et al., 2019; Turco et al., 2019; Vargas et al., 2019). TANK-binding kinase 1 (TBK1) is known to interact with and phosphorylate these adaptor proteins, increasing their affinity for both ULK1/FIP200 complex components and for ubiquitin (Moore and Holzbaur, 2016; Pilli et al., 2012; Richter et al., 2016; Vargas et al., 2019; Zachari et al., 2019).

Ubiquitination involves the sequential action of three classes of enzymes: ubiquitin activating enzymes (E1); ubiquitin conjugating enzymes (E2); and ubiquitin ligation enzymes (E3). While the human genome encodes only two E1 and ∼40 E2 enzymes, it encodes more than 600 E3 enzymes. It is the E3 ligases that determine ubiquitination substrate selectivity. The best-studied E3 ligase involved in mitophagy is the Parkinson disease-associated protein Parkin, which is activated following mitochondrial uncoupling (Nguyen et al., 2016a; Pickrell and Youle, 2015). However, additional E3 ligases are also reported to contribute to mitophagy in response to other stimuli (Chen et al., 2017; Zachari et al., 2019). In general, these ligases are primarily thought to ubiquitinate the damaged mitochondria and thus play a role in mitophagy-substrate recognition. In principle, E3 ligases could also directly regulate the mitophagy machinery, but this concept has received less attention in the literature.

The <u>tri</u>partite <u>m</u>otif (TRIM) protein family consists of more than 80 genes in humans, most of which encode a RING ubiquitin ligase domain at their N-terminus (Short and Cox, 2006). A remarkably high percentage of TRIM proteins have been shown to function as mediators of selective autophagy and to interact with autophagy proteins (Di Rienzo et al., 2020; Hatakeyama, 2017). In most cases, the role of the TRIM’s ubiquitin ligase activity in autophagy has not been addressed. We recently demonstrated that TRIM5α, a protein best known for roles in antiviral defense (Cloherty et al., 2021), can localize to mitochondria where it plays an essential role in ubiquitin-dependent mitophagy (Saha et al., 2022). We reported that TRIM5α’s activity as an ubiquitin ligase is required for its mitophagy functions. However, TRIM5α was not required for the bulk ubiquitination of mitochondria following damage; suggesting that the role of TRIM5α-mediated ubiquitination in mitophagy differs from that of Parkin or other E3 ligases that ubiquitinate mitochondrial proteins.

Here, we carried out an in-depth mechanistic dissection of TRIM5α-mediated mitophagy and identified two overlapping activities. First, mitochondrial damage induced TRIM5α-mediated K63-linked poly-ubiquitination of TBK1 and itself. This resulted in the formation of a multi-protein scaffold including TRIM5α, TBK1, and mitophagy adaptors that promotes the activation of TBK1 on mitochondria. This TRIM5α-ubiquitin-TBK1 axis was required for the downstream recruitment of the ULK1/FIP200 complex to damaged mitochondria. Second, TRIM5α directly recruits ULK1 to damaged mitochondria through protein-protein interactions that are ubiquitin-independent. Our studies indicate that additional TRIMs may utilize similar mechanisms to promote mitophagy. Collectively, these findings position TRIMs at a central hub in orchestrating mitophagy and establish an upstream TRIM-ubiquitin-TBK1 axis in mitophagy that may also be applicable to other forms of TRIM-mediated selective autophagy.

## RESULTS

### TRIM5α and TBK1 interact in response to mitochondrial damage and TRIM5α recruits active TBK1 to mitochondria

We previously showed that TRIM5α was responsible for two distinct ubiquitin-dependent mitophagy pathways: Parkin-dependent mitophagy induced by CCCP and for Parkin-independent mitophagy induced by ivermectin (Saha et al., 2022). Proximity ligation-based proteomic analysis indicated a possible interaction between TRIM5α and TBK1 (Saha et al., 2022). Since TBK1 is a required factor for both CCCP- and ivermectin-induced mitophagy pathways (Heo et al., 2015; Zachari et al., 2019), we hypothesized that TRIM5α might engage TBK1 in mitophagy. To test this concept, we first determined whether mitochondrial damaging agents could increase TRIM5α-TBK1 interactions. Under basal conditions in transfected HEK293T cells, GFP-tagged TRIM5α immunoprecipitated myc-TBK1 (Figure 1A, B). Mitophagy inducers CCCP (Fig 1A) or ivermectin (Fig 1B) both reproducibly increased the abundance of TBK1 in protein complexes with GFP-TRIM5α. When active, TBK1 is phosphorylated at serine 172 (pTBK1). We found that pTBK1 co-immunoprecipitated with TRIM5α, indicating that TRIM5α interacts with active TBK1 (Figure 1A, B). Time course experiments showed that interactions between GFP-TRIM5α and myc-TBK1 increased within the first 30 minutes following mitochondrial uncoupling with CCCP (Figure 1C). Coincidently with the increased TRIM5α-TBK1 interaction, we also observed the appearance of high molecular weight (>250 kDa) bands detected with anti-GFP antibody, indicative of post-translational modifications of TRIM5α in response to mitochondrial damage. Stably expressed HA-tagged TRIM5α colocalized with endogenous pTBK1 in ivermectin-treated HeLa cells (Figure 1D). TRIM5α and pTBK1 colocalization was found to be closely associated with mCherry-Parkin signal on mitochondria following CCCP treatment (Figure S1A). Proximity ligation (PLA) can be used to identify protein-protein interactions within 40 nm *in situ*, with positive PLA signal being detected as a fluorescent punctum. We observed numerous PLA signals indicating TRIM5α-TBK1 interactions in CCCP-treated cells (Figure 1E). Most, but not all, of these PLA signals were found to colocalize with Parkin-decorated mitochondria, indicating that TRIM5α-TBK1 complexes form on the surface of damaged mitochondria.

**Figure 1.**
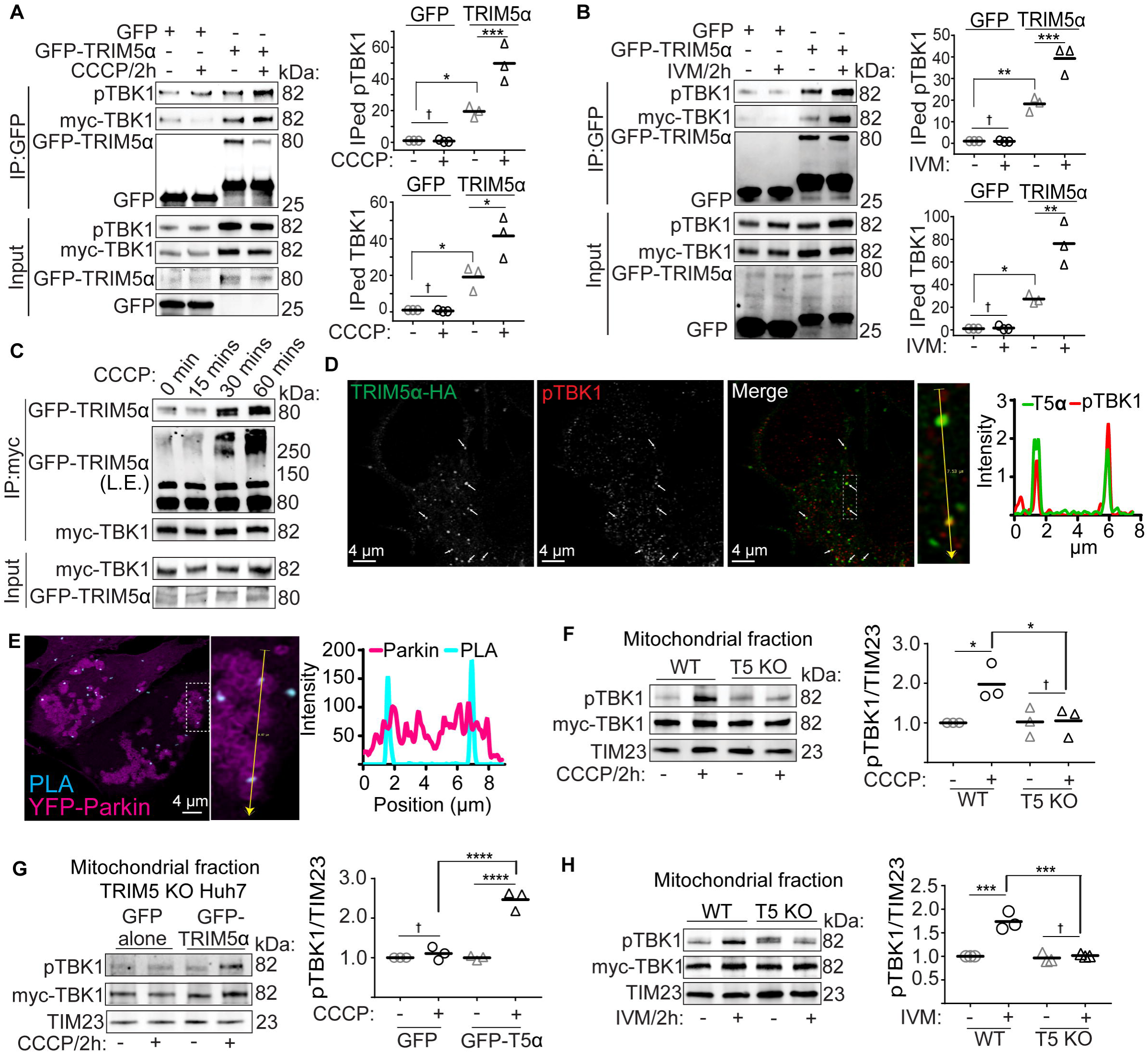
TRIM5α interacts with TBK1 and promotes its localization to damaged mitochondria. **(A, B)** Coimmunoprecipitation analysis of interactions between TRIM5α and TBK1 under basal conditions or 2 hours after treatment with mitochondrial damaging agents CCCP (A) or invermectin (IVM) (B) in transiently transfected HEK293T cells. Plots show the relative abundance of immunoprecipitated (IPed) total TBK1 or phosphorylated TBK1 (pSer172) relative to the abundance of IPed GFP or GFP-TRIM5α. **(C)** Coimmunoprecipitation analysis of the kinetics of TRIM5α-TBK1 interactions following CCCP treatment in transiently transfected HEK293T cells. L.E., long-exposure. **(D)** Confocal microscopic analysis of TRIM5α and pTBK1 localization in HeLa cells stably expressing HA-tagged TRIM5α following 1 h treatment with IVM. Arrows indicate colocalized puncta. The zoomed-in imaged with the intensity profile is from the boxed region. **(E)** Proximity ligation assay (PLA, pseudo-colored cyan) analysis of TRIM5α-HA interactions with pTBK1 in cells expressing YFP-Parkin (pseudo-colored magenta). PLA dots indicate sites where TRIM5α and pTBK1 are within 40 nm of each other. Intensity profile of the boxed region is shown at the right. **(F)** The abundance of total and phosphorylated TBK1 in mitochondrial fractions from WT or TRIM5 knockout Huh7 cells treated or not with CCCP for 2 h. Plot shows the relative abundance of pTBK1 normalized to the mitochondrial protein TIM23. Each data point represents an independent experiment. **(G)** TRIM5 KO Huh7 cells were transiently transfected with GFP-TRIM5α or GFP alone and treated with DMSO or CCCP for 2 h prior to isolation of mitochondrial fractions and immunoblotting as shown. Quantitation of the relative abundance of pTBK1, normalized to TIM23, is shown at the right. **(H)** Immunoblot analysis of mitochondrial fractions obtained from WT and TRIM5 KO Huh7 cells treated or not with IVM for two hours. Plot shows the relative abundance of pTBK1, normalized to TIM23, from three independent experiments. Data: mean + SEM; *P* values determined by ANOVA with Tukey’s multiple comparison test; *, *P* < 0.05; **, *P* < 0.01; ***, *P* < 0.001; ****, *P* < 0.0001; †, not significant.

We next tested whether TRIM5α was important for recruiting or retaining active pTBK1 to mitochondria following mitochondrial damage. In WT Huh7 cells transfected with myc-TBK1, either CCCP or ivermectin treatment increases the abundance of pTBK1 in mitochondrial fractions (Figure 1F, H). In contrast, pTBK1 was not enriched in mitochondrial fractions from cells lacking the TRIM5 gene (Figure 1F, H). Transient expression of GFP-TRIM5α in TRIM5 knockout cells restored the CCCP-induced pTBK1 accumulation on damaged mitochondria (Figure 1G). In these experiments, TRIM5α primarily impacted the mitochondrial abundance of active pTBK1 rather than total TBK1. Consistent with these results, pTBK1 showed reduced colocalization with the mitochondrial marker TIM23 in TRIM5 knockout Huh7 cells relative to control Huh7 cells under CCCP-treatment conditions (Figure S1B). Together, these data show a requirement for TRIM5α in localizing active TBK1 to mitochondria following the induction of Parkin-dependent and Parkin-independent mitophagy.

### TBK1 is required for TRIM5α-mediated recruitment of autophagy adaptors and regulators to damaged mitochondria

We tested whether TRIM5α knockout influenced the phosphorylation of TBK1 substrates. The protein Sequestosome-1 (SQSTM1, also p62) is recruited to ubiquitinated mitochondria (Wong and Holzbaur, 2014) and is phosphorylated at serine 349 (pSQSTM1) by TBK1 (Richter et al., 2016). By high content imaging, we observed a robust increase in the number of pSQSTM1 puncta following either CCCP or ivermectin treatment in wild type Huh7 cells that was substantially abrogated in TRIM5 knockout cells (Figure 2A, B).

**Figure 2.**
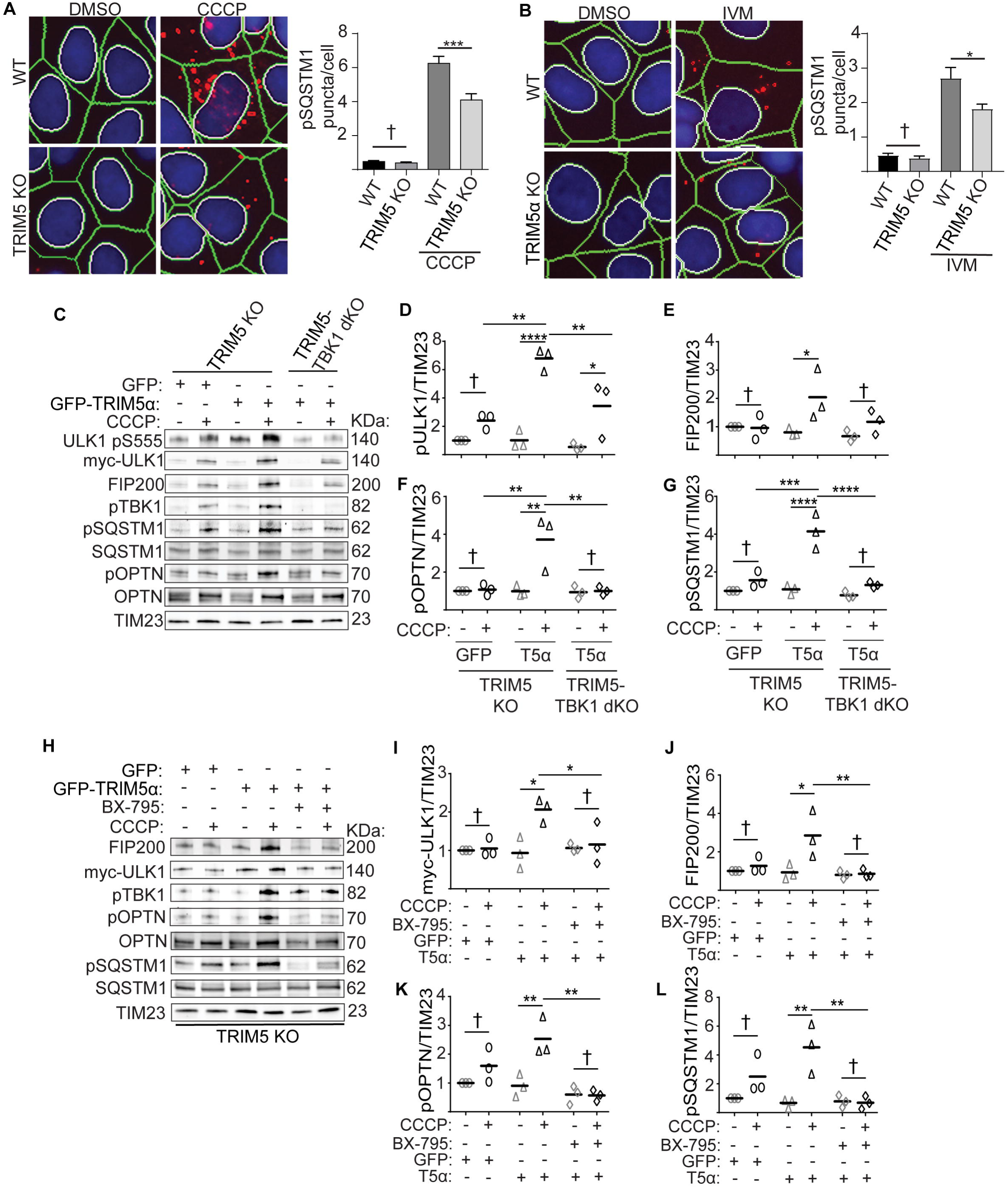
TRIM5α recruits autophagy machinery to damaged mitochondria in a TBK1-dependent manner. **(A, B)** The number of phospho-SQSTM1 (pSer349) puncta was determined in WT and TRIM5 KO Huh7 cells under basal conditions or following 2 h treatment with CCCP (A) or IVM (B) by high content imaging. **(C-G)** TRIM5 knockout or TRIM5 and TBK1 double knockout (dKO) Huh7 cells were transfected with GFP-TRIM5α (T5α) or GFP alone as indicated and treated or not with CCCP (4 h) prior to isolation of mitochondrial fractions and immunoblotting (C). The abundance of the indicated proteins was determined, normalized to TIM23 as a loading control, and plotted (D-G). **(H-L)** TRIM5 KO Huh7 cells were transfected with GFP-TRIM5α (T5α) or GFP alone and treated or not with CCCP for 4 hours in the presence or absence of the TBK1 inhibitor BX-795 prior to isolation of mitochondrial fractions and immunoblotting. The results of three independent experiments are quantitated in I-L. TIM23 was used as a loading control. Data: mean + SEM; *P* values determined by ANOVA with Tukey’s multiple comparison test; *, *P* < 0.05; **, *P* < 0.01; ***, *P* < 0.001; ****, *P* < 0.0001; †, not significant.

We next tested whether TBK1 was downstream of TRIM5α in mitophagy. We previously showed that the recruitment of several autophagy proteins, including members of the ULK1/FIP200 complex, to damaged mitochondria is dependent on TRIM5α (Saha et al., 2022). Ectopic expression of TRIM5α in TRIM5 knockout cells rescues this phenotype (Figure S2A). We generated TRIM5-TBK1 double knockout cells (Figure S2B) to test whether this effect is dependent on TBK1. As expected, expression of GFP-TRIM5α in TRIM5 knockout Huh7 cells increased the CCCP-driven recruitment of total and autophagy-active ULK1 (phosphorylated at Ser555) and of FIP200 to mitochondria (Figure 2C-E). TRIM5α re-expression also enhanced the abundance of TBK1-phosphorylated autophagy adaptors SQSTM1 (phospho-Ser349) and optineurin (phospho-Ser177) in mitochondrial fractions from CCCP-treated cells (Figure 2C-G). However, the absence of TBK1 completely abrogated the TRIM5α-dependent localization of pULK1, FIP200, pSQSTM1, and pOptineurin to mitochondria in response to CCCP. Similarly, treatment with the TBK1 inhibitor BX-795 also prevents the TRIM5α-dependent localization of mitophagy proteins to CCCP-damaged mitochondria, supporting a role for TBK1 downstream of TRIM5α in mitophagy (Figure 2H-L).

In addition to TBK1, TRIM5α has also been shown to activate the kinase TAK1 (Fletcher et al., 2018). We used the TAK1 inhibitor (5Z)-7-Oxozeaenol to test whether TAK1 contributed to CCCP-induced mitophagy, as measured by the degradation of the mitochondrial inner membrane protein COXII, but saw no effect (Figure S2C, D). Our results establish TBK1 as an essential factor downstream of TRIM5α in mitophagy.

In addition to their actions in mitophagy, both TRIM5α and TBK1 have well-known roles in antiviral defense. Ectopic expression of TRIM5α in HEK293T cells was sufficient to enhance the abundance of pTBK1 in cells under both basal conditions and following CCCP treatment, possibly through stabilization of total TBK1 (Figure S2E-G). TRIM5α expression also increased the abundance of pSQSTM1 (Figure S2E, H), an autophagy-relevant TBK1 substrate, but only following CCCP treatment. Given this result, we asked whether TRIM5α expression could promote the TBK1-dependent activation of interferon signaling using a dual luciferase reporter system. As previously reported (Fletcher et al., 2018), TRIM5α expression induces activation of NF-κB driven luciferase (Figure S2I). However, and despite its ability to promote TBK1 activation as measured by pTBK1 abundance, TRIM5α expression actually reduced the ability of TBK1 to promote activation of an interferon-stimulated response element (Figure S2J). These data indicate that the TRIM5α-activated pool of TBK1 does not promote innate immune signaling and, instead, is specific to autophagy.

### TRIM5α-mediated K63-linked ubiquitination of TBK1 is important for mitophagy

Previous reports have shown that K63-linked ubiquitination of TBK1 is important for its activity in interferon signaling (Song et al., 2016; Tu et al., 2013). We hypothesized that TRIM5α’s ubiquitin ligase activity, which we previously showed to be important for mitophagy (Saha et al., 2022), could promote mitophagy through TBK1 ubiquitination. Expression of GFP-TRIM5α in HEK293T cells robustly increased the appearance of high molecular weight TBK1 and the ability of TBK1 to coimmunoprecipitate ubiquitin relative to cells expressing GFP alone (Figure 3A), indicative of TBK1 ubiquitination. Remarkably, both CCCP and ivermectin triggered TBK1 ubiquitination that was seen as early as 1 hour after treatment and continued to increase up to at least 4 hours after treatment. In these experiments, mitochondrial damage-induced TBK1 ubiquitination was almost entirely dependent on TRIM5 (Figure 3B and Figure S3A). We saw similar results when we monitored TBK1 ubiquitination by an ubiquitin mutant that was disabled for forming all linkages except for K63, showing that TRIM5α promotes activating (K63-linked) ubiquitination of TBK1 in response to mitochondrial damage (Figure 3C). Since TBK1 ubiquitination has not previously been described in response to conditions that induce mitophagy or other types of autophagy, we tested whether other known inducers of autophagy, namely amino acid starvation, chemical inhibition of mTOR with pp242, or endomembrane damage caused by LLOMe, could also trigger this posttranslational modification of TBK1 similarly to mitochondrial damage with CCCP. However, of these, CCCP was the only treatment that induced TBK1 ubiquitination (Figure S3B).

**Figure 3.**
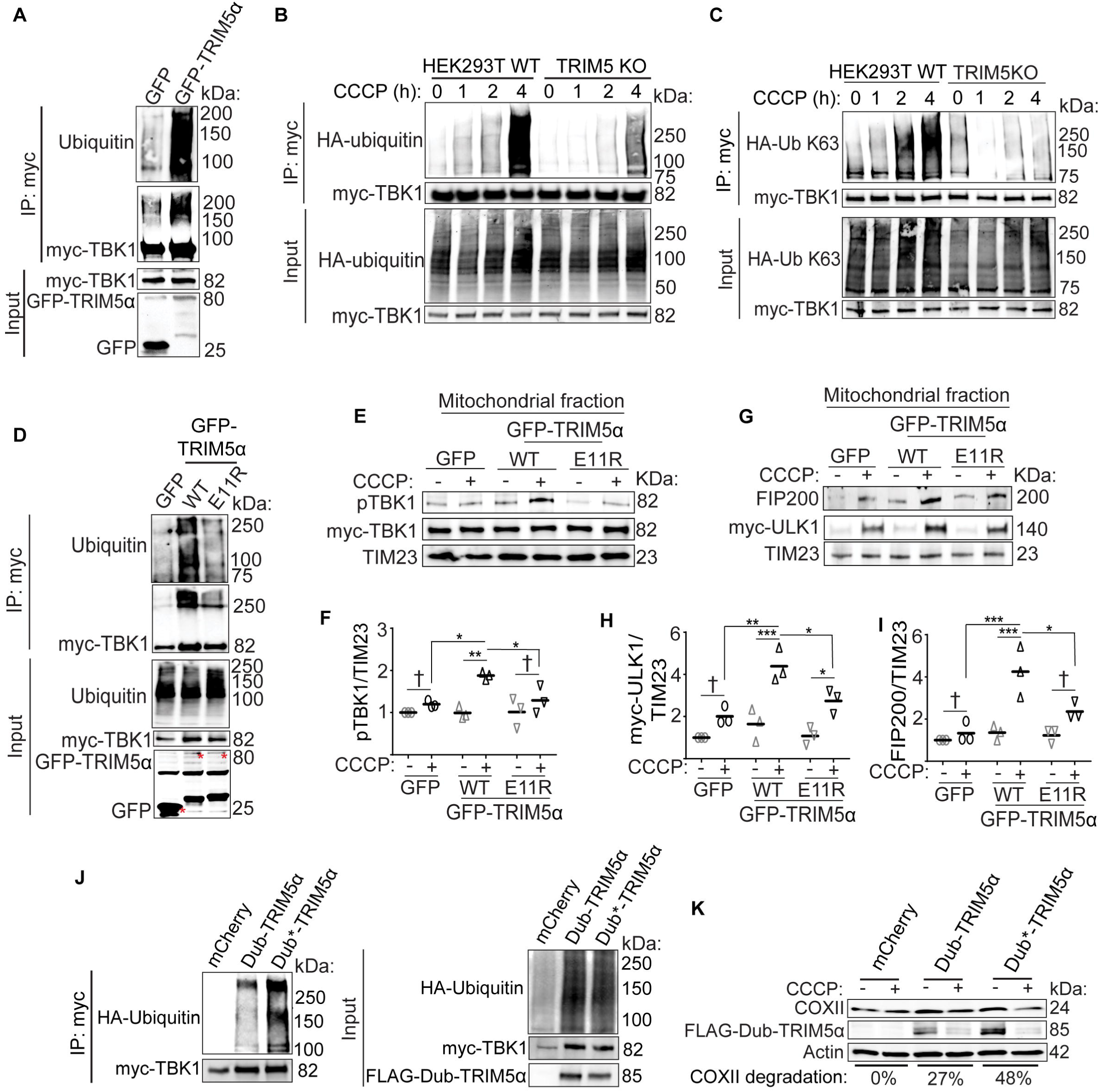
TRIM5α’s ubiquitin ligase activity is required for K63-linked poly-ubiquitination of TBK1 and for the recruitment of autophagy proteins to damaged mitochondria. **(A)** Coimmunoprecipitation analysis of TBK1 ubiquitination in HEK293T cells transfected with myc-TBK1 and GFP alone or GFP-TRIM5α. **(B)** The effect of CCCP treatment on TBK1 ubiquitination in WT and TRIM5 knockout HEK293T cells. **(C)** The effect of CCCP treatment on the K63-linked poly-ubiquitination of TBK1 in WT or TRIM5 KO HEK293T cells. Cells were transfected with myc-TBK1 and HA-tagged ubiquitin that is only capable of generating K63 linkages (HA-UB K63). **(D)** Coimmunoprecipitation analysis of TBK1 ubiquitination in HEK293T cells expressing GFP alone or GFP-tagged WT or E11R TRIM5α. Red * indicate GFP or GFP-TRIM5α bands of the expected molecular weight. **(E, F)** Mitochondrial fractions were obtained from TRIM5 knockout Huh7 cells that had been transiently transfected with myc-TBK1 and GFP or GFP-tagged WT or E11R TRIM5α and treated or not with CCCP for 2 h. The abundance of pTBK1 in the mitochondrial fractions from three independent experiments was determined, normalized to TIM23, and plotted in (F). **(G-I)** The abundance of the indicated proteins in mitochondrial fractions isolated from TRIM5 knockout Huh7 cells transfected with myc-ULK1 and GFP alone or GFP-tagged TRIM5α (WT or E11R) and treated or not with CCCP. The relative abundance of myc-ULK1 or FIP200 from three independent experiments is shown in panels H and I, respectively. **(J)** Coimmunoprecipitation analysis of TBK1 ubiquitination in HEK293T cells transfected with myc-TBK1 and TRIM5α fused to either a catalytically active (Dub) or catalytically inactive (Dub*) deubiquitinase. mCherry was used as a negative control. **(K)** Relative COXII degradation following CCCP treatment in TRIM5 knockout Huh7 cells transfected with FLAG-Dub-TRIM5α, FLAG-Dub*-TRIM5α, or mCherry alone and treated or not with CCCP for 18 h. *P* values determined by ANOVA with Tukey’s multiple comparison test; *, *P* < 0.05; **, *P* < 0.01; ***, *P* < 0.001; ****, *P* < 0.0001; †, not significant.

We next tested whether TRIM5α’s enzymatic activity as an E3 ligase is important for its ability to recruit mitophagy proteins to mitochondria in response to damage. Replacing glutamine 11 in TRIM5α with arginine (E11R) disrupts the ability of TRIM5α to interact with E2 conjugating enzymes and interferes with TRIM5α’s enzymatic activity without disrupting its overall structure (Fletcher et al., 2018). Unlike what we see with wild type TRIM5α, expression of TRIM5α E11R does not increase TBK1 ubiquitination (Figure 3D), demonstrating that TRIM5α directly catalyzes TBK1 ubiquitination. TRIM5α E11R was also significantly attenuated in its ability to restore the recruitment of active pTBK1 (Figure 3E, F) and both ULK1 and FIP200 (Figure 3G-I) into mitochondrial fractions following CCCP treatment of TRIM5 knockout cells. Nevertheless, neither E11R nor another E2-binding patch mutant of TRIM5α (L19R) were different from WT in their ability to stabilize TBK1 in whole cell lysates (Figure S3C-E). Together, these results suggest that TRIM5α-mediated TBK1 ubiquitination is a prerequisite step for the mitochondrial localization of active TBK1 in response to mitochondrial damage, but is not required for the overall stabilization of TBK1.

As an alternative approach to assess the importance of TRIM5α-mediated TBK1 ubiquitination, we tested whether the enzymatic removal of ubiquitin from TBK1 (or other TRIM5α-interacting proteins) interfered with mitophagy. This was accomplished using a TRIM5α fusion protein that links the catalytic domain of the UL36 deubiquitinase from herpes simplex virus to TRIM5α’s N terminus (Dub-TRIM5α) (Campbell et al., 2016). As a negative control for these experiments, we also employed a deubiquitinase-dead TRIM5α fusion (Dub*-TRIM5α). While Dub-TRIM5α still somewhat increased TBK1 ubiquitination relative to cells transfected with a control protein (mCherry), it did so to a much lesser extent than did Dub*-TRIM5α (Figure 3J). Consistent with this, Dub-TRIM5α failed to restore CCCP-induced COXII degradation in TRIM5-knockout Huh7 cells (Figure 3K). In contrast, Dub*-TRIM5α, which is a surrogate for wild type TRIM5α because it has intact ubiquitination function, potently restored mitophagy in these experiments (Figure 3K). In summary, our data reveal that TRIM5α mediates ubiquitination of TBK1, particularly under conditions of mitochondrial damage. Furthermore, abrogation of TRIM5α-dependent ubiquitination of TBK1 (or other substrates) impairs TRIM5α’s ability to mediate the recruitment of autophagy proteins to damaged mitochondria and to execute mitophagy.

### TRIM5α ubiquitinates TBK1 at K30 and K401 to enhance TBK1 interactions with autophagy adaptors NDP52, SQSTM1, and NBR1

Structural and functional studies of TBK1 have demonstrated the importance of TBK1 ubiquitination at lysine residues 30 and 401 for full TBK1 activity in cells (Song et al., 2016; Tu et al., 2013). We mutated both of these sites in TBK1, replacing them with arginine (TBK1 2X K→R), to test whether these are the predominant ubiquitination sites of TBK1 following mitochondrial damage. As shown in Figure 4A, WT TBK1 showed a basal level of K63-linked poly-ubiquitination that strongly and progressively increased after CCCP treatment. However, the 2X K→R mutant of TBK1 displayed markedly reduced K63-linked ubiquitination in untreated cells and no increase in ubiquitination one hour following CCCP treatment. At 2 hours post CCCP treatment, the 2X K→R TBK1 showed a dramatic increase in ubiquitination with comparable ubiquitination to WT, indicating that additional sites on TBK1 are eventually ubiquitinated under conditions of mitochondrial damage. We next tested whether TRIM5α ubiquitinates TBK1 at K30 and K401 (Figure 4B). Whereas co-expression of GFP-TRIM5α with wild type myc-TBK1 massively increased K63-linked TBK1 ubiquitination, ubiquitination of the 2X K→R mutant of TBK1 was reduced by ∼75%. However, TRIM5α clearly promotes ubiquitination of TBK1 at additional sites, since we still observed considerable TRIM5α-dependent ubiquitination of 2X K→R TBK1.

**Figure 4.**
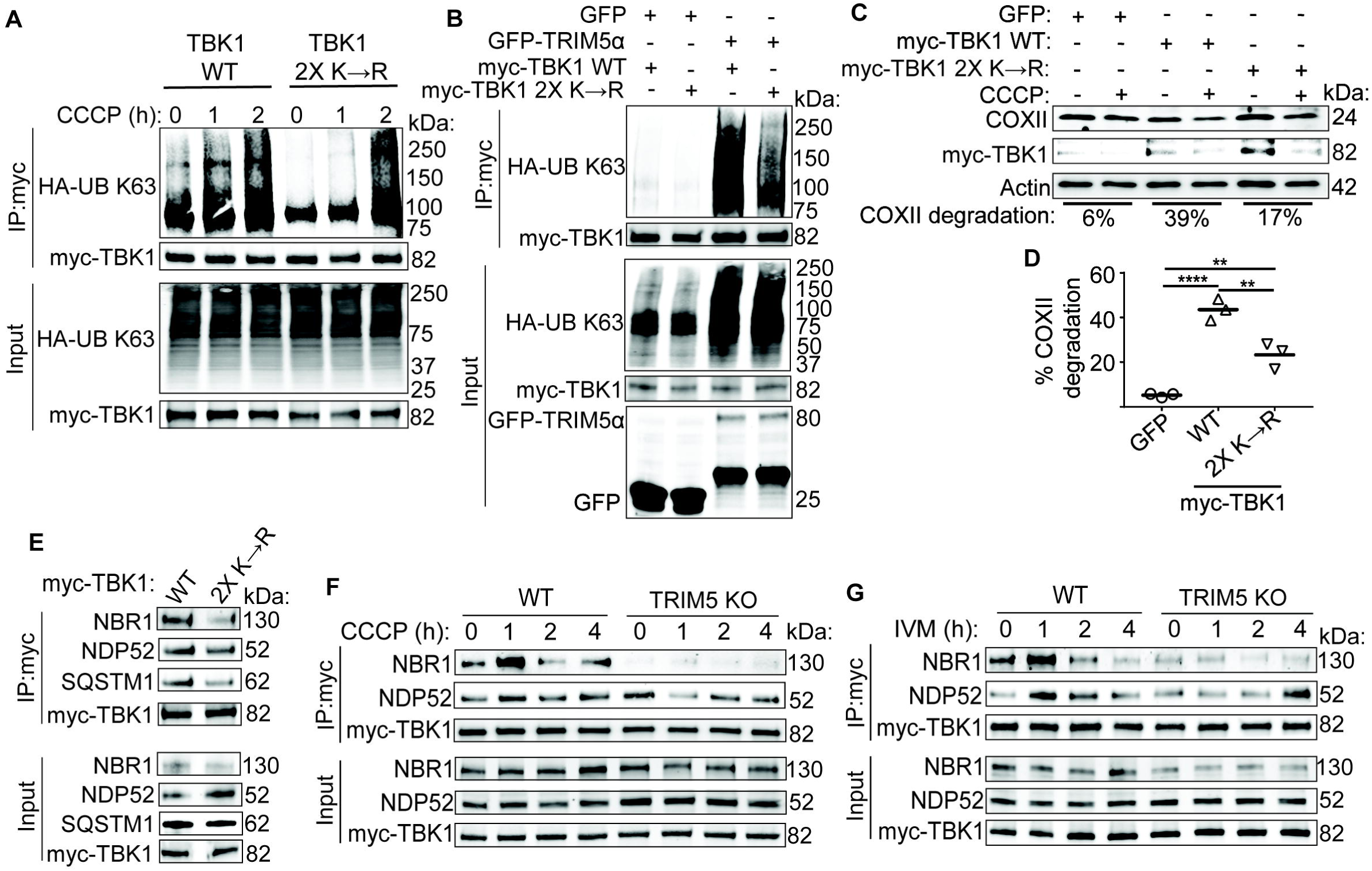
Ubiquitination of TBK1 at K30 and K401 by TRIM5α is required for mitophagy and enhances TBK1 interactions with autophagy adaptors. **(A)** Coimmunoprecipitation analysis of TBK1 ubiquitination in HEK293T cells transfected with WT or K30R/K401R (2X K→R) mutant myc-TBK1, and K63-restricted HA-tagged ubiquitin (HA-UB K63), followed by treatment with CCCP for the indicated time points. **(B)** TBK1 ubiquitination assay in HEK293T TRIM5KO cells cotransfected with plasmids encoding HA-UB K63 and either the control vector GFP or GFP-TRIM5α and WT myc-TBK1 or 2X K→R mutant myc-TBK1. **(C, D)** The ability of WT or 2X K→R mutant TBK1 to rescue CCCP-induced mitophagy in TRIM5 KO Huh7 cells. GFP was used as a negative control. The relative extent of CCCP-induced COXII degradation from three independent experiments is plotted in (D). **(E)** Coimmunoprecipitation analysis of interactions between WT or 2X K→R TBK1 mutant and endogenous autophagy adaptors NBR1, NDP52, and SQSTM1 from transiently transfected HEK293T cells. **(F, G)** The impact of TRIM5 knockout on CCCP (F) or IVM (G) induced interactions between myc-TBK1 and endogenous NDP52 or NBR1. HEK293T cells were transfected with the indicated plasmid and treated with mitophagy inducers prior to lysis and immunoprecipitation with anti-myc. *P* values determined by ANOVA with Tukey’s multiple comparison test; **, *P* < 0.01; ***, *P* < 0.001; ****, *P* < 0.0001.

Our data demonstrate that K30 and K401 are the major sites of TRIM5α-dependent TBK1 ubiquitination, and that these sites are ubiquitinated in response to mitochondrial damage. This suggests a model in which K63-linked poly-ubiquitination of TBK1 at K30 and K401 may be important for mitophagy. To test this idea, we measured the CCCP-induced degradation of the mitochondrial protein COXII in TBK1 knockout Huh7 cells transfected with either WT or 2X K→R TBK1 or an irrelevant protein (GFP) as a negative control (Figure 4C, D). In the GFP-transfected cells, we observed minimal COXII degradation, in line with the fact that TBK1 is essential for CCCP-induced mitophagy. Expression of WT TBK1 in these cells significantly increased COXII degradation in response to CCCP. However, expression of the 2X K→R TBK1 mutant was significantly less efficient at promoting CCCP-induced COXII degradation despite roughly comparable levels of TBK1 protein expression (Figure 4C, D). These data demonstrate that ubiquitination of TBK1 at K30 and K401 is important for TBK1’s mitophagy functions.

One possible way in which ubiquitination could increase TBK1 function is through facilitating the recruitment of TBK1 co-factors that could mediate proper TBK1 localization or activation. TBK1 interacts with a set of five autophagy adaptors (SQSTM1, NDP52, Optineurin, TAX1BP1, and NBR1) (Zellner et al., 2021), all of which encode ubiquitin-binding domains. TBK1 ubiquitination by TRIM5α could promote these interactions and/or could contribute to the assembly of TBK1-adaptor clusters held together by avidity-based interactions. Accordingly, WT TBK1 more efficiently coimmunoprecipitates NBR1, NDP52, and SQSTM1 than does the ubiquitination deficient 2X K→R mutant of TBK1 (Figure 4E). We then tested whether mitochondrial damage with either CCCP or ivermectin, both of which induce TBK1 ubiquitination (Figure 3 and S3), could impact protein-protein interactions between TBK1 and autophagy adaptors, using NDP52 and NBR1 as representatives. We found that both treatments transiently increased the ability of myc-TBK1 to coimmunoprecipitate endogenous NDP52 and NBR1 (Figure 4F, G). This effect was lost in TRIM5 knockout cells. Intriguingly, while TBK1 ubiquitination continues to increase up to four hours after treatment, TBK1 interactions with autophagy adaptors peaked at one hour. The reasons for this kinetic discrepancy require further study. Overall, these findings indicate that TRIM5α-dependent K63-linked ubiquitination of TBK1 is important for the formation of TBK1-adaptor protein complexes, possibly explaining the importance of this post-translational modification to TBK1-dependent mitophagy.

### TRIM5α interacts with TBK1 through K63-linked poly-ubiquitin/autophagy adaptor intermediates

In domain mapping experiments, we found that the TRIM5α RING domain was required for TRIM5α-TBK1 interactions in cells (Figure S4A, B). However, GST-pulldown experiments failed to show any interaction between recombinant TBK1 and either full-length TRIM5α or the N terminal portion of TRIM5α (Figure S4C), indicating that TRIM5α-TBK1 interactions might require additional factors. Since the RING domain encompasses TRIM5α’s ubiquitin ligase activity and is the site of TRIM5α auto-ubiquitination, it is possible that TRIM5α-mediated ubiquitination could contribute to TRIM5α-TBK1 interactions. In support of this concept, we observed that mitochondrial damaging agents increased the appearance of high molecular weight TRIM5α simultaneously with TRIM5α-TBK1 binding (Figure 1C). Both ivermectin and CCCP treatment dramatically increased TRIM5α ubiquitination in a time-dependent manner as measured by increases in the amount of ubiquitin coimmunoprecipitating with TRIM5α and the appearance of high molecular weight TRIM5α bands, the latter of which could represent either ubiquitinated TRIM5α and/or its organization into stabilized higher-order structures (Figure 5A and S4D). Follow-up experiments demonstrated that TRIM5α was subject to K63-linked ubiquitination in response to ivermectin and CCCP (Figure S4E, F). We found that the ivermectin-induced K63-linked poly-ubiquitination of TRIM5α is largely dependent on TRIM5α’s ubiquitin ligase activity, since the E11R mutant of TRIM5α showed much less ubiquitination than did wild type (Figure 5B). This result demonstrates that TRIM5α is primarily auto-ubiquitinated, and that its ability to carry out auto-ubiquitination is enhanced following mitochondrial damage. However, we still observed ivermectin-induced K63-linked ubiquitination of TRIM5α E11R at 2 h after treatment, implicating additional ubiquitin ligases as secondary contributors to TRIM5α ubiquitination in response to mitochondrial damage (Figure 5B).

**Figure 5.**
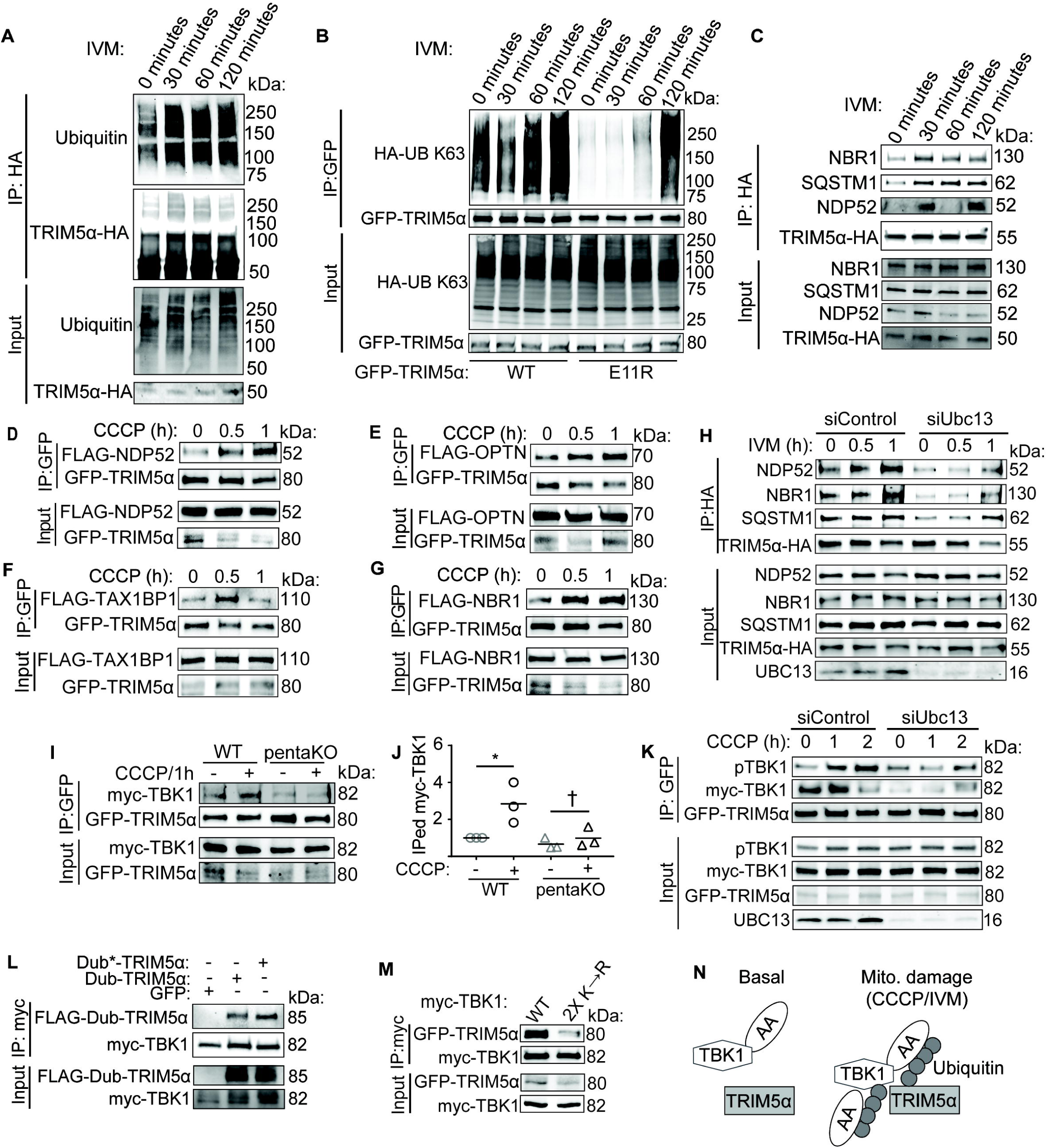
Interactions between TRIM5α and TBK1 are mediated by ubiquitin-binding autophagy receptors. **(A)** Coimmunoprecipitation analysis of TRIM5α ubiquitination in HeLa cells stably expressing TRIM5α-HA and treated with IVM for the indicated time points. **(B)** Analysis of K63-linked ubiquitination of WT or E11R TRIM5α following IVM treatment of transfected TRIM5α knockout HEK293T cells. **(C)** HeLa cells expressing TRIM5α-HA were treated with IVM for the indicated time points prior to lysis, immunoprecipitation with anti-HA, and immunoblotting. **(D-G)** Coimmunoprecipitation analysis of interactions between GFP-TRIM5α and FLAG-tagged autophagy receptors NDP52, OPTN, TAX1BP1, and NBR1 from lysates of transiently transfected HEK293T cells treated with CCCP for the indicated time. **(H)** Coimmunoprecipitation analysis of interactions between TRIM5α and the indicated autophagy adaptors in HeLa cells stably expressing HA-tagged TRIM5α that are transfected with control siRNA (siControl) or UBC13 siRNA for 48 h and then treated with IVM for the indicated time periods. Lysates were subjected to immunoprecipitation with anti-HA and immunoblots probed as indicated. **(I, J)** Coimmunoprecipitation analysis of interactions between GFP-TRIM5α and myc-TBK1 in HeLa cells lacking all five autophagy adaptors (SQSTM1, NBR1, NDP52, OPTN, and TAX1BP1; pentaKO) or parental HeLa cells treated or not with CCCP for 1 h. Both WT and pentaKO cells were also transfected with mCherry-Parkin. The abundance of immunoprecipitated (IPed) myc-TBK1 was determined from three independent experiments, normalized to IPed GFP-TRIM5α, and plotted in (H). *P* values determined by ANOVA with Tukey’s multiple comparison test; *, *P* < 0.05; †, not significant. **(K)** The impact of Ubc13 knockdown on interactions between GFP-TRIM5α and myc-TBK1 in HEK293T cells treated with CCCP for the indicated time as determined by co-immunoprecipitation with anti-GFP antibodies. **(L)** Lysates from HEK293T cells transfected with myc-TBK1 and TRIM5α fused to an active deubiquitinase (Dub) or a catalytically inactive deubiquitinase (Dub*) with an N-terminal FLAG tag were subjected to immunoprecipitation with anti-myc prior to immunoblotting as shown. GFP was used as a negative control. **(M)** Coimmunoprecipitation analysis of interaction between GFP-TRIM5α and WT or 2X K→R mutant myc-TBK1 in HEK293T TRIM5 KO cells. **(N)** Proposed mechanism for mitochondrial-damaged induced TRIM5α-TBK1 interactions. The assembly of these complexes likely occurs on the surface of damaged mitochondria. AA, autophagy adaptor (e.g. NDP52, SQSTM1, etc.)

The increase in TRIM5α ubiquitination was mirrored by an increase in the interactions between TRIM5α and autophagy adaptors. As shown in Figure 5C, ivermectin treatment increased the ability of TRIM5α to coimmunoprecipitate endogenous NBR1, SQSTM1, and NDP52 within 30 minutes. The interaction between TRIM5α and SQSTM1 or NBR1 remained at high levels for at least the next 90 minutes; while with NDP52 interactions were dynamic, reproducibly showing strong binding at 30 minutes and variable binding at later time points. In transiently transfected HEK293T cells, CCCP treatment also increased the interactions between TRIM5α and NDP52, optineurin, TAX1BP1, and NBR1 (Figure 5D-G) but not SQSTM1 (Figure S4G). We found that the interactions between TRIM5α and autophagy adaptors required ubiquitination, as knockdown of the E2 conjugating enzyme Ubc13/Ube2n, which specifically mediates the formation of K63-linked poly-ubiquitin chains, abrogates the ivermectin-induced interaction between TRIM5α and endogenous NDP52, NBR1, and SQSTM1 (Figure 5H). These data reveal previously unrecognized, ubiquitin-dependent, interactions between TRIM5α and autophagy adaptors that are responsive to mitochondrial damage.

We hypothesized that TRIM5α-TBK1 interactions could be mediated indirectly through shared interactions with autophagy adaptors. To test this, we determined whether TRIM5α and TBK1 could interact in cells lacking all five autophagy adaptors (PentaKO cells) (Lazarou et al., 2015). As expected based on our prior experiments, both CCCP and ivermectin increased TRIM5α-TBK1 coimmunoprecipitation in WT HeLa cells. However, this increase was not seen in the PentaKO HeLa cells (Figure 5I, J and S4H). This result demonstrates that one or more autophagy adaptors, all of which interact with both TRIM5α and TBK1 (Zellner et al., 2021), can scaffold TRIM5α-TBK1 interactions. Given the role of TBK1 ubiquitination in its interactions with a subset of autophagy receptors (Figure 4), the importance of K63-linked ubiquitination in TRIM5α’s interactions with autophagy adaptors (Figure 5H), and the importance of autophagy adaptors in mediating TRIM5α-TBK1 interactions (Figure 5I, J), we next tested the hypothesis that ubiquitination contributes to TRIM5α-TBK1 interactions. Accordingly, knockdown of Ubc13 prevented the time-dependent interactions between TRIM5α and both TBK1 and pTBK1 following mitochondrial damage with CCCP (Figure 5K). Deubiquitinase-fused TRIM5α was also markedly less efficient at interacting with TBK1 than was deubiquitinase-dead fused TRIM5α (Figure 5L), further emphasizing the importance of ubiquitination in TRIM5α-TBK1 interactions. Finally, we found that ubiquitinated residues K30 and K401 in TBK1 are also important contributors to TRIM5α-TBK1 interactions (Figure 5M), consistent with these residues’ role in allowing interactions between TBK1 and autophagy adaptors (Figure 4E).

Our model for how TRIM5α and TBK1 interact in response to mitochondrial damage is shown in Figure 5N. Either CCCP or ivermectin enhance TRIM5α-mediated K63-linked ubiquitination of itself and of TBK1. Ubiquitin chains attached to either TRIM5α or TBK1 or both then attract autophagy adaptors, which can then further bridge TRIM5α and TBK1 through shared interactions. In this way, TRIM5α can promote the condensation, concentration, and activation of TBK1 and autophagy adaptors on the mitochondria through self-amplifying multivalent interactions, thus laying a foundation for the recruitment of the autophagy machinery.

### The role of the TRIM5 LC3-interacting region (LIR) in mitophagy

We and others have previously shown that TRIM5α encodes two LC3 interacting regions (LIR) (Keown et al., 2018; Mandell et al., 2014). LIR motifs mediate interactions with autophagy proteins, most notably with LC3 family proteins (LC3A, LC3B, and LC3C) and their homologues (GABARAP, GABARAP L1 and GABARAP L2), collectively referred to as mammalian Atg8 proteins (mAtg8s) (Birgisdottir et al., 2013). Mutation of the second of these LIR motifs (LIR2; Figure 6A) is sufficient to reduce TRIM5α interactions with mAtg8s without disrupting TRIM5α-mediated retroviral capsid-specific restriction (Keown et al., 2018), a phenotype that requires the maintenance of TRIM5α’s overall three-dimensional structure. We next compared the ability of wild type and LIR2 mutant TRIM5α to restore CCCP-induced mitophagy in TRIM5 knockout cells. As seen in other experiments, wild type TRIM5α substantially increased CCCP-induced COXII degradation (Figure 6B, C). However, the expression of LIR2 mutant TRIM5α had no effect, showing that an intact LIR motif is required for TRIM5α -directed mitophagy.

**Figure 6.**
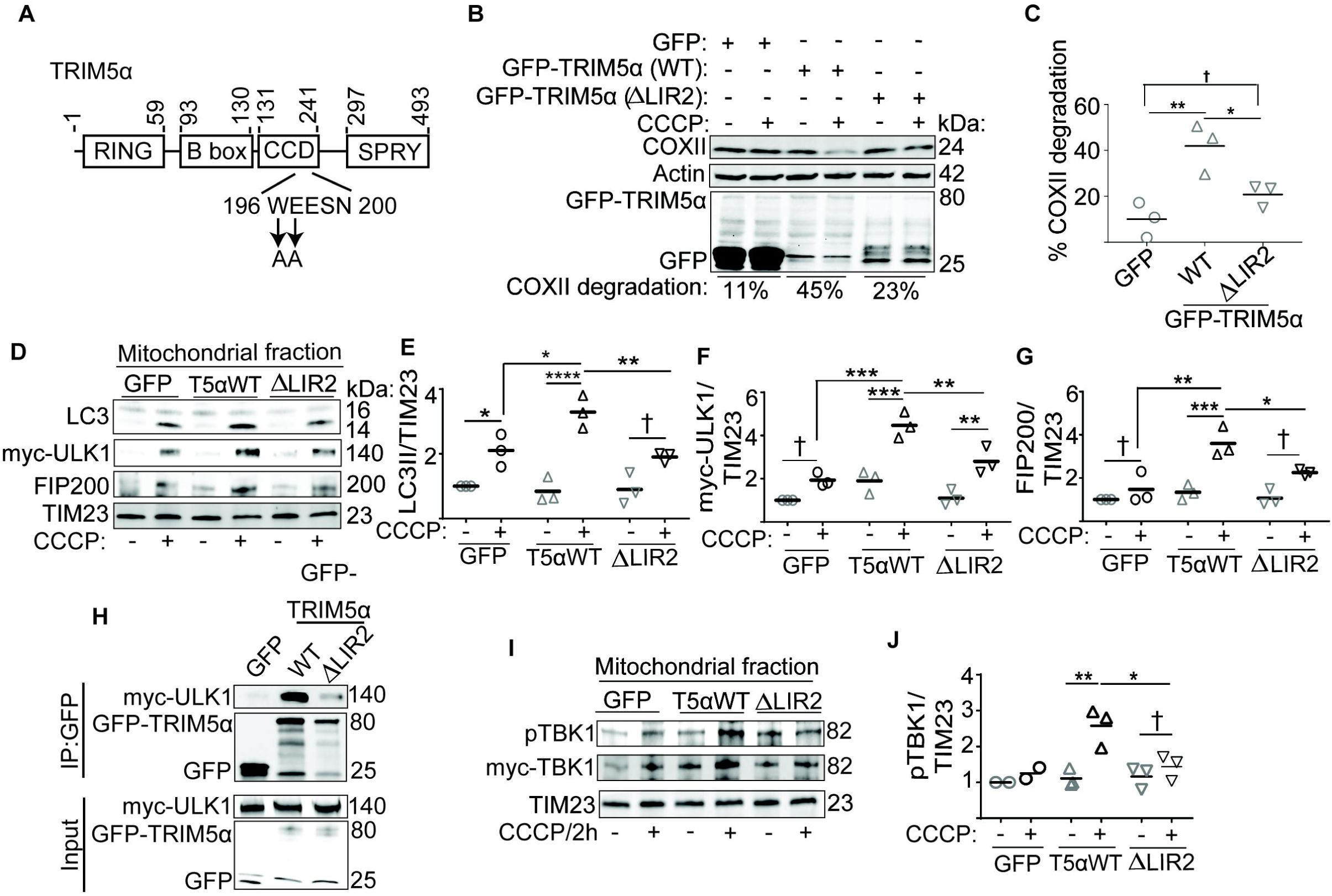
Role of the TRIM5α LIR motif in mitophagy. **(A)** Schematic of TRIM5 domain structure and the location of the TRIM5α LIR motif mutated in this study. **(B, C)** Immunoblot analysis of CCCP-induced COXII degradation in TRIM5 knockout Huh7 cells complemented with WT or LIR2-mutant GFP-TRIM5α or GFP alone. The relative abundance of COXII was determined, normalized to actin, and the results of three independent experiments plotted in C. **(D-G)** Immunoblot analysis of proteins in mitochondrial fractions isolated from TRIM5 knockout Huh7 cells transfected with myc-ULK1 and GFP-TRIM5α (WT or ΔLIR2) or GFP prior to treatment with CCCP or DMSO for 4 hours. The abundance of the indicated proteins was determined, normalized to TIM23 and plotted in E-G. **(H)** Coimmunoprecipitation analysis of myc-tagged ULK1 in protein complexes with GFP-tagged TRIM5α (WT or ΔLIR2) or GFP alone. Lysates from transiently transfected HEK293T cells were immunoprecipitated with anti-GFP and subjected to immunoblotting as shown. **(I, J)** Immunoblot analysis of proteins in mitochondrial fractions isolated from TRIM5 knockout Huh7 cells transfected with myc-TBK1 and GFP-TRIM5α (WT or ΔLIR2) or GFP prior to treatment with CCCP or DMSO for 2 hours. The abundance of pTBK1 was determined, normalized to TIM23 and plotted in J. *P* values determined by ANOVA with Tukey’s multiple comparison test; *, *P* < 0.05; **, *P* < 0.01; ***, *P* < 0.001; ****, *P* < 0.0001; †, not significant.

Mitochondrial fractionation experiments with TRIM5 knockout cells expressing GFP alone or GFP-tagged wild type- or LIR2 mutant-TRIM5α demonstrated that expression of wild type TRIM5α increased the abundance of autophagy active (lipidated) LC3B associated with mitochondria, while this was not the case with the LIR2 mutant (Figure 6D, E). Unexpectedly, the LIR2 mutant TRIM5α was also less effective than WT in promoting the recruitment of ULK1 and FIP200 (Figure 6D, F, and G), suggesting an expanded role for TRIM5α’s LIR in mitophagy beyond interactions with mAtg8s. LIR motifs have recently been shown to bind to FIP200 (Turco et al., 2019), raising the possibility that TRIM5α’s LIR motifs could be involved in assembling TRIM5α complexes with ULK1/FIP200 complexes. While mutation of LIR2 in TRIM5α did not affect TRIM5α’s interaction with either FIP200 or ATG13 (another component of the ULK1/FIP200 complex; Figure S5A, B), mutation of TRIM5α’s LIR2 motif dramatically reduced the ability of GFP-TRIM5α to coimmunoprecipitate myc-ULK1 (Figure 6H) and for myc-ULK1 to coimmunoprecipitate GFP-TRIM5α (Figure S5C). TRIM5α interactions with ULK1 are not mediated by mAtg8s, since deletion of all six mAtg8s (Nguyen et al., 2016b) does not abrogate TRIM5α-ULK1 coimmunoprecipitation (Figure S5D). TRIM5α’s LIR2 motif was also not required for TRIM5α-TBK1 interactions (Figure S5E) or for the ability of TRIM5α over-expression to stabilize TBK1 (Figure S5F). Nevertheless, LIR2 mutant TRIM5α was attenuated in its ability to restore the localization of active pTBK1 to damaged mitochondria in TRIM5α knockout cells relative to what is seen with wild type TRIM5α (Figure 6I, J). Together, these data suggest that the role of TRIM5α’s LIR motif is to mediate TRIM5α-ULK1 interactions, and that these interactions are important for mitophagy.

### TRIM5α cooperates with TRIM27 in executing mitophagy

In addition to TRIM5α’s role in ubiquitin-dependent mitophagy, we recently showed that another TRIM, TRIM27 (also known as Ret finger protein or RFP), contributes to ubiquitin-independent mitophagy mediated by FKBP8 (Garcia-Garcia et al., 2023) and promotes the mitochondrial localization of pTBK1, albeit through unknown mechanisms. TRIM5α and TRIM27 are closely related genes, both members of the Class IV sub-family of TRIMs (Short and Cox, 2006). Sequence analysis shows that TRIM27 has a putative LIR motif that aligns with TRIM5’s LIR2 but does not retain TRIM5’s LIR1 (Figure S6A). A tiled TRIM27 peptide array reveals that this motif, defined by the sequence 184-WEFEQL-189, is essential for binding to *in vitro* translated GABARAP (Figure S6B). GST pull-down analysis showed that TRIM27 directly interacts with the mAtg8s LC3A, LC3C, and all three GABARAPs (Figure S6C), results similar to those reported for TRIM5α (Mandell et al., 2014). Substitution of TRIM27 amino acids 184, 186, and 189 with alanine substantially abrogated TRIM27’s ability to interact with mAtg8s (Figure S6D). Together, these data establish TRIM27 as a mAtg8 interactor with a strong preference for LC3C and demonstrate that these interactions are mediated by a *bona fide* LIR motif.

The similarities between TRIM27 and TRIM5α made us ask whether these TRIMs employ overlapping mechanisms in mitophagy. In support of this concept, we found that ectopic expression of GFP-TRIM27 was just as efficient at restoring mitophagy in TRIM5 knockout Huh7 cells as was expression of GFP-TRIM5α (Figure 7A). Proteomic analysis of TRIM5α interacting partners under basal conditions revealed TRIM27 as weak hit (Saha et al., 2022), and we confirmed that TRIM5α and TRIM27 interact by coimmunoprecipitation experiments in transfected HEK293T cells (Figure 7B). TRIM5α-HA and GFP-TRIM27 colocalize in cytoplasmic structures observed by confocal microscopy in HeLa cells (Figure S6E). Importantly, TRIM5α/TRIM27-double positive puncta localized to discreet sub-domains of mCherry-Parkin-positive mitochondria (Figure 7C), positioning them to have similar functions in mitophagy. Ivermectin treatment increased the formation of TRIM5α-TRIM27 protein complexes (Figure 7D), showing that TRIM27, like TBK1 and autophagy adaptors, is recruited into TRIM5α complexes under mitophagy-inducing conditions. A previous study has shown direct interaction between TRIM27 and TBK1 (Zheng et al., 2015), suggesting that TRIM27 may facilitate TRIM5α-dependent mitophagy by helping to assemble TRIM5α-TBK1 complexes. Indeed, TRIM5α-HA was better able to coimmunoprecipitate myc-TBK1 in invermectin-treated cells expressing GFP-TRIM27 than in ivermectin-treated cells expressing GFP-alone (Figure 7E). Together, these studies demonstrate that TRIM27 promotes TRIM5α- and ubiquitin-dependent mitophagy, likely through actions focused on TBK1. In conclusion, our studies with TRIM5α and TRIM27 provide a model for how TRIM-directed mitophagy functions.

**Figure 7.**
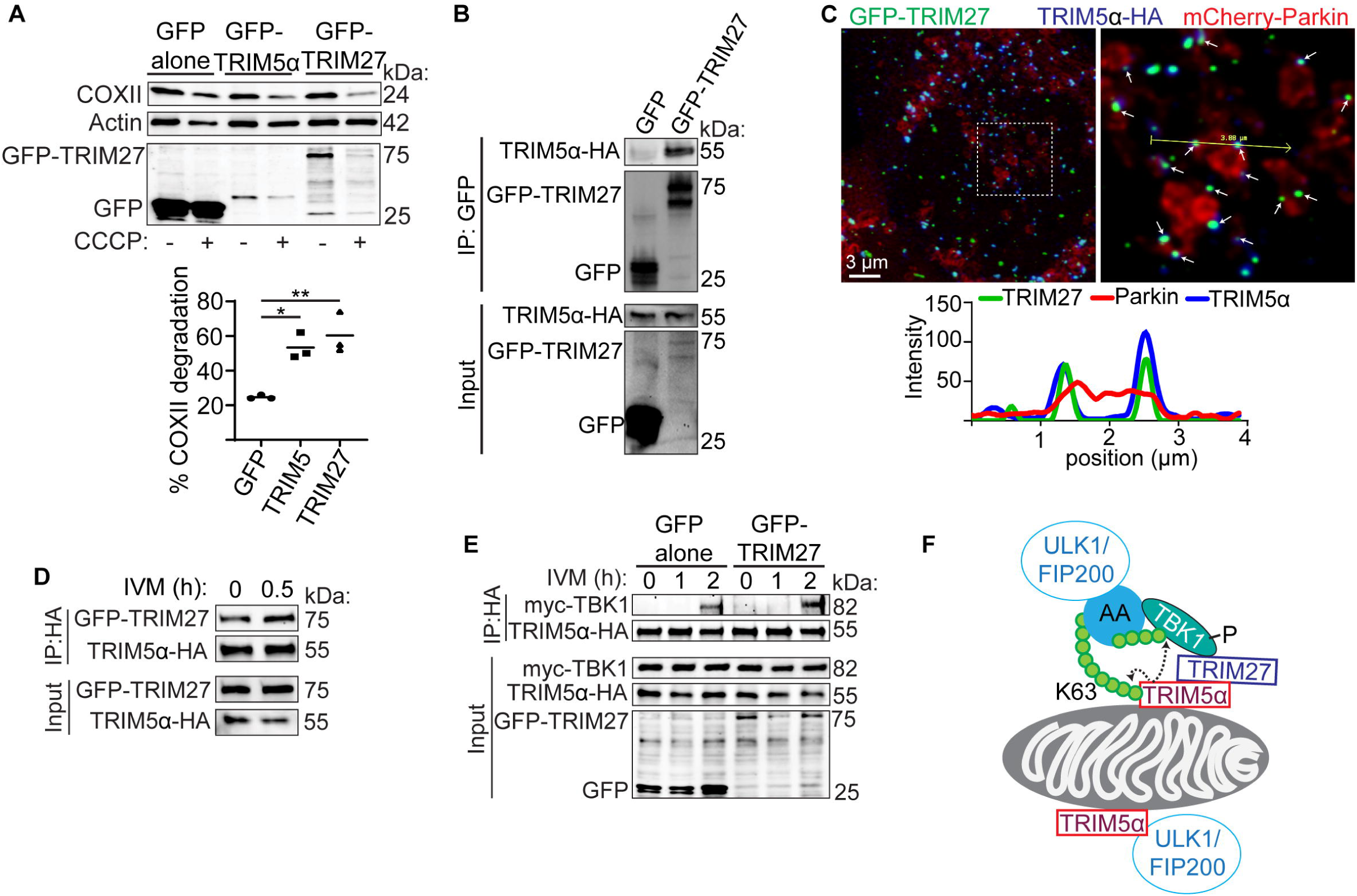
TRIM27 interacts with TRIM5α and can restore mitophagy in TRIM5 knockout cells. **(A)** Immunoblot analysis of COXII protein abundance in TRIM5 knockout Huh7 cells transfected with GFP-TRIM5α, GFP-TRIM27, or GFP alone and treated or not with CCCP for 18 h. Plot shows the relative extent of COXII degradation following CCCP treatment for three independent experiments. **(B)** Coimmunoprecipitation analysis of interactions between GFP alone or GFP-TRIM27 and HA-tagged TRIM5α in HeLa cells. **(C)** HeLa cells stably expressing TRIM5α-HA were transfected with GFP-TRIM27 and mCherry-Parkin and then treated with CCCP for 90 m prior to fixation and confocal microscopy. The boxed in region (left) is shown on the right. White arrows indicate structures positive for TRIM5α and TRIM27 signal that are adjacent to coalescent Parkin signal. The intensity profile for the yellow line is shown to the right. **(D)** Coimmunoprecipitation analysis of TRIM5α and TRIM27 interactions in HeLa cells treated with IVM for the indicated time points. **(E)** Lysates from HeLa cells stably expressing TRIM5α-HA and transfected with myc-TBK1 and either GFP-TRIM27 or GFP alone and treated with IVM for the indicated time were subjected to immunoprecipitation with anti-HA and immunoblotted as indicated. **(F)** Proposed dual roles for TRIM5α in mitophagy. Top, TRIM5α indirectly recruits the upstream autophagy regulatory machinery (ULK1, FIP200, etc.) through a K63-linked ubiquitin/autophagy adaptor/TBK1 axis involving TRIM27. TRIM5α’s enzymatic activity as an E3 ubiquitin ligase is required for this mode of TRIM5α-mediated mitophagy. AA, autophagy adaptor. Bottom, TRIM5α recruits the ULK1 complex through direct interactions involving its LC3-interacting region (LIR2). *P* values determined by ANOVA with Tukey’s multiple comparison test; *, *P* < 0.05; **, *P* < 0.01.

## DISCUSSION

The molecular events leading up to mitophagy involve the concerted recruitment of multiple individual proteins and protein complexes to the mitochondria, often as part of feed-forward loops that rapidly amplify the local concentration of mitophagy factors and induce their activation. Our study shows that TRIM5α, and potentially other TRIMs like TRIM27, contributes to these events in two ways. The first way requires TRIM5α’s enzymatic activity as an E3 ubiquitin ligase (Figure 7F, top). Mitochondrial stress induces the K63-linked poly-ubiquitination of TRIM5α. These ubiquitin chains serve as scaffolds that link TRIM5α with ubiquitin-binding autophagy adaptor proteins (e.g. NDP52, Optineurin, p62/SQSTM1, NBR1, and TAX1BP1). All of these adaptor proteins also interact with TBK1, and thus they serve as intermediaries that bridge TRIM5α-TBK1 interactions. This allows TRIM5α to catalyze K63-linked ubiquitination of TBK1 at K30 and K401, promoting TBK1 activity while also providing additional landing sites for autophagy adaptor recruitment and subsequent assembly of TRIM5α-ubiquitin-TBK1 complexes in a self-amplifying manner.

This TRIM-ubiquitin-TBK1 axis is required for the proper activation and localization of TBK1 in mitophagy. K63-linked ubiquitination of TBK1 is essential for TBK1 activity in immune signaling (Song et al., 2016; Tu et al., 2013), although this concept has received less attention with regards to TBK1’s actions in selective autophagy. Our data show that this post-translational modification of TBK1 is increased in response to mitochondrial damage in a TRIM5α-dependent manner. Active TBK1 can phosphorylate autophagy adaptors near their ubiquitin-binding domains, promoting the adaptors’ ability to interact with ubiquitinated TRIM5α, TBK1, or mitochondrial proteins (Richter et al., 2016). Phosphorylation of autophagy adaptors near their LIR motifs by TBK1 enhances the adaptors’ ability to interact with and recruit mAtg8 proteins and/or the ULK1/FIP200 complex (Vargas et al., 2019; Wild et al., 2011). TBK1 can also promote mitophagy through phosphorylation of RAB7A (Heo et al., 2018) or by directly activating the phosphatidylinositol 3-kinase (PI3K) complex I (Nguyen et al., 2023). By activating TBK1, TRIM5α is upstream of all of these processes.

A prerequisite for the actions of TBK1 in mitophagy is that TBK1 must be localized to damaged mitochondria and activated. The mitochondrial recruitment of TBK1 can be mediated by autophagy adaptors. We found that TRIM5α-dependent ubiquitination is required for the enhanced interactions between TBK1 and a subset of autophagy adaptors in response to mitochondrial damaging agents. We presume that this is a result of TRIM5α promoting TBK1 ubiquitination, since the 2X K→R mutant of TBK1 is both less ubiquitinated by TRIM5α and shows reduced interactions with autophagy adaptors. By promoting interactions between autophagy adaptors and TBK1, TRIM5α enables TBK1’s recruitment to damaged mitochondria.

In many respects, the downstream outcomes of TRIM5α’s ubiquitin-dependent mitophagy actions mirror the roles of Parkin or other E3 ligases previously implicated in mitophagy. However, the primary difference is the nature of the ubiquitinated targets. In Parkin-dependent mitophagy, Parkin ubiquitinates a wide variety of proteins on the mitochondrial outer membrane (Sarraf et al., 2013). In contrast, TRIM5α appears to ubiquitinate itself and TBK1. Since our prior work showed that a pool of TRIM5α localizes to mitochondria even under basal conditions (Saha et al., 2022), this allows for the assembly of TRIM5α-ubiquitin-TBK1 complexes where they are needed when mitochondria are damaged.

Our data show that TRIM5α may not be unique among TRIMs in its mitophagy-directed actions on TBK1. TRIM27 shows substantial homology to TRIM5α, and we previously reported that TRIM27 contributes to an ubiquitin-independent form of mitophagy (Garcia-Garcia et al., 2023). Here, we showed that TRIM27 expression can rescue mitophagy in TRIM5 knockout cells. This, along with the strong colocalization of TRIM5α and TRIM27 on the surface of mitochondria following mitochondrial damage suggests that TRIM5α and TRIM27 operate through overlapping mechanisms to execute mitophagy. However, the finding that TRIM27 could enhance TRIM5α interactions with TBK1 in response to mitochondrial damage suggests that these two proteins may not be fully redundant. More study is necessary to address whether TRIM27 uses the same mechanism(s) as does TRIM5α in mitophagy or to determine if there are cell type- or context-dependent differences between mitophagy mediated by these two TRIMs.

The second mode by which TRIM5α contributes to mitophagy is through direct interactions with the autophagy machinery (Figure 7F, bottom). We found that mutation of an LC3-interacting (LIR) motif in TRIM5α abrogated interactions between ULK1 and TRIM5α in coimmunoprecipitation experiments. TRIM5α LIR mutants were similar to wild type in terms of their interactions with FIP200, ATG13, and TBK1, and the loss of LC3 or GABARAP proteins (mAtg8s) did not impact TRIM5α/ULK1 interactions. This suggests that mutation of the LIR motif of TRIM5α likely directly impacts the ability of TRIM5α and ULK1 to interact. Accordingly, LIR-mutant TRIM5α was attenuated in its ability to enhance the CCCP-dependent mitochondrial recruitment of ULK1 in TRIM5 knockout cells, suggesting that TRIM5α could directly recruit ULK1 to damaged mitochondria through protein-protein interactions. Interestingly, although no reproducible difference was seen between WT and LIR2-mutant TRIM5α’s ability to coimmunoprecipitate TBK1, the LIR mutant of TRIM5α was also attenuated in its ability to recruit active TBK1 to damaged mitochondria. This unexpected result could be explained by the finding that ULK1’s ubiquitin-independent recruitment to mitochondria can precede and promote Parkin-dependent actions such as recruitment of TBK1 to damaged mitochondria (Hung et al., 2021). Whether TRIM5α contributes to the rapid and transient recruitment of ULK1 to damaged mitochondria is a subject for future study.

In conclusion, our study positions TRIMs at a central node in mitophagy regulation (Figure S6F), from which they can orchestrate mitophagy through direct protein-protein interactions or through regulating the activation and localization of TBK1. In addition to TRIM5α and TRIM27, many other TRIMs have been implicated in regulating selective autophagy of a variety of substrates (Di Rienzo et al., 2020). The TRIM-ubiquitin-TBK1 axis provides a new lens through which the mechanistic basis for selective autophagy by other TRIMs could be viewed.

### Limitations of the study

In this study, we determined the mechanisms of TRIM5α-mediated mitophagy in immortalized cancer cell lines. We found that chemical agents that damage mitochondria and induce mitophagy increase TRIM5α-mediated ubiquitination of itself and of TBK1, but we have not identified how mitochondrial damage is translated into activation of TRIM5α’s enzymatic activity. Presumably, this involves the higher-order assembly of TRIM5α molecules as is seen by TRIM5α in the context of retroviral capsid recognition (Fletcher et al., 2018) or by TRIM72 in response to recognition of vesicle structures (Park et al., 2023). Using a broad array of coimmunoprecipitation assays, we established that TRIM5α and TBK1 likely interact indirectly in a manner requiring K63-linked poly-ubiquitin chains and autophagy adaptors, but efforts to predict how these interactions form on a structural level using AlphaFold or structural docking software (PDBePISA) failed to yield high-confidence models. We also did not determine the relative importance of the five autophagy adaptors in mediating TRIM5α/TBK1 interactions. Finally, although our model implicates TBK1 ubiquitination by TRIM5α in scaffolding interactions between TBK1 and autophagy adaptors, we noticed that TBK1/adaptor interactions peaked at early time points after mitophagy-inducing treatments while TBK1 ubiquitination continued to increase. Experiments to explain this kinetic discrepancy are beyond the scope of this study.

## Acknowledgements

This work was supported by P20GM121176 and R01AI155746 to M.A.M from the US National Institutes of Health. We thank Dr. Ed Campbell (Loyola University Chicago) for sharing reagents the TRIM5α deubiquitinase fusion plasmids and Drs. Richard Youle and Michael Lazarou (US NIH and Monash University, respectively) for sharing the PentaKO and HexaKO cells. Dr. Jing Pu (University of New Mexico) commented on the manuscript.

## Author contributions

B.S., G.L.W., S.O., G.E., H.O., and M.A.M performed research; B.S., H.O., E.S., and M.A.M designed research and analyzed data; B.S. and M.A.M. wrote the paper. The authors declare that they have no conflict of interest.

## MATERIALS AND METHODS

### Cell culture

HEK293T, HeLa, and Huh7 cells were obtained from the American Type Culture Collection (ATCC) and grown in Dulbecco’s modified Eagle’s medium (Life Technologies,11965126) supplemented with 10% fetal bovine serum (FBS, Life technologies, 26140-079), 100 U/ml penicillin and 100 µg/ml streptomycin at 37°C in a 5% CO2 atmosphere. HeLa cells stably expressing HA-tagged HuTRIM5α were obtained from Joseph Sodroski (Harvard) and were maintained in the above media supplemented with 1 µg/ml puromycin. Generation of TRIM5 knockout HEK293T, HeLa, and Huh7 cells using CRISPR/Cas9 based gene editing were described previously (Saha et al., 2022) and were maintained in the above media supplemented with 200 µg/ml hygromycin. WT and mitophagy adaptor pentaKO cells (gifts from Dr. Richard J. Youle, National Institutes of Health, USA), WT and mATG8 HexaKO cells (gifts from Dr. Michael Lazarou, Monash University, Australia) were cultured in the same manner. TBK1 knockout HEK293T and TRIM5-TBK1 double knockout Huh7 cells were generated by transduction with lentiCRISPRv2-based lentiviruses followed by 2-4 weeks of culturing in medium containing 8 µg/ml blasticidin. Knockout lines were confirmed by immunoblot.

### Generation of knockout cell lines using CRISPR/cas9 gene editing

Viral particles for the generation of knockout cell lines were produced by transfecting HEK293T cells with a lentiviral vector, lentiCRISPRv2 carrying both Cas9 enzyme and a guide RNA targeting specific gene together with the packaging plasmids psPAX2 and pMD2.G at the ratio of 10 µg, 10 µg and 10 µg/10 cm dish. Transfections were carried out by using ProFection Mammalian Transfection System (Promega, E1200), medium was changed 16h post transfection and virus containing supernatant was harvested 48h later, clarified by centrifuging for 5 min at 1200 rpm, 0.45 µm-filtered (Millipore, SE1M003M00), diluted with full medium at 1:1 ratio and used to transduce target cells for 48 h.

### Cloning and transfection

GFP-TRIM5α and myc-TBK1 were mutated using a site-directed mutagenesis kit (Agilent, 210518). All plasmid constructs generated in this study were validated by DNA sequencing. All other TRIM5α mutants used in this study were described earlier (Mandell et al., 2014). Myc-TBK1 and FLAG-tagged autophagy adaptor plasmids were gift from Dr. Vojo Deretic. The AP1 luciferase reporter plasmid was a gift from Alexander Dent (Addgene plasmid #40342; 3XAP1pGL3), the NF-κB luciferase reporter was purchased from Promega (#E8491), the Renilla luciferase plasmid (pRL-SV40, Addgene plasmid #27163) was a gift from Ron Prywes and ISRE luciferase plasmid was gifted by Dr. Michael Gale (University of Washington). Dub-TRIM5α (rhesus) and Dub*-TRIM5α were a gift from Dr. Edward Campbell. . pDest-EGFP-TRIM27 was described earlier (Garcia-Garcia et al., 2023). Myc-TRIM27 W184A/F186A/L189A was mutated with Quickchange II site-directed mutagenesis kit (Agilent). pDONR221-AZI2/NAP1 was synthesized by Invitrogen GeneArt services (Thermo Fisher Scientific). pDest15-AZI2/NAP1 was made with Gateway LR cloning (Thermo Fisher Scientific). Plasmid transfections were performed using Lipofectamine 2000 (ThermoFisher, 11668019) or Calcium Phosphate (Promega, E1200). Samples were prepared for analysis the day after DNA transfection.

All siRNA smart pools were from Dharmacon. siRNA was delivered to cells using Lipofectamine RNAiMAX (ThermoFisher, 13778150). For siRNA experiments, cells were harvested 48–72h after siRNA transfection.

### Treatments

Working concentrations for reagents were as follows: CCCP, 10 µM for overnight experiments; and 20 µM for experiments ≤ 4 hours; IVM, 15 µM; BX-795, 10 µM; 5Z-7-Oxozeaenol, 10 µM; pp242, 10 µM; LLOMe, 100 µM; MG132, 10 µM.

### Western blotting, immunoprecipitation, and immunofluorescent labeling

For immunoprecipitation experiments, cells seeded in 10 cm dishes were transfected with specific constructs to overexpress proteins of interest for 24 h, followed by the indicated treatments in the presence of MG132. Cells were then lysed using ice cold lysis buffer (150 mM Tris-buffered Saline, 50 mM NaCl, with 0.5% v/v Triton-X 100) supplemented with EDTA-free cOmplete protease inhibitor, PMSF, and phosphatase inhibitors. Samples were incubated on ice for 30 min. Beads were equilibrated using the lysis buffer mentioned above. Protein lysates were precleared by centrifugation at 4°C for 15 min at 17,000 g. Clarified lysates were incubated with specific equilibrated beads for various periods at 4°C. Beads were then washed with ice cold Wash buffer (150 mM Tris-buffered Saline, 50 mM NaCl) 3 to 4 times. Bound proteins were eluted with LDS lysis buffer with 50 mM DTT. Samples were boiled for an additional 10 min. For ubiquitination related immunoprecipitation experiments, a deubiquitinase inhibitor (PR-619; 10 µM, UBPBio) was added in lysis buffer and in wash buffers in addition to the protease inhibitors detailed above. For these experiments, we lysed cells in RIPA buffer (50 mm Tris, 0.1% SDS, 0.5% deoxycholate, 1% Nonidet P-40, 150 mM NaCl) to reduce non-covalent interactions between proteins and ubiquitin.

SDS PAGE was carried out using pre-cast poly-acrylamide gels and immunoblotting was performed using standard procedures (Mandell et al., 2014). Immunoblot data was acquired using a Chemidoc MP instrument (Biorad) and quantitatively analyzed using Biorad Image Lab software.

For immunoflourescent labeling of samples for high content imaging or confocal experiments, samples were fixed in 4% paraformaldehyde (Sigma) for 30 minutes prior to permeabilization in buffer containing 0.1% Saponin (Sigma) and 3% BSA. Following 1 hour incubation in primary antibodies and extensive washing with phosphate buffered saline (PBS), AlexaFluor-conjugated secondary antibodies (Life Technologies) were used. Coverslips were mounted in ProLong Diamond anti-fade reagent (Life Technologies).

### Confocal microscopy

Sub-airy unit (0.6AU) pinhole confocal microscopy with a Leica TCS-SP8 microscope was performed followed by computational image restoration with Huygens Essential (Scientific Volume Imaging, Hilversum, Netherlands) utilizing a constrained maximum likelihood estimation algorithm. All 3D images were acquired with a 63X/1.4NA plan apochromat oil immersion objective and sampled at ideal Nyquist sampling rates in x, y, and z planes. Voxel lateral and axial dimensions were determined by utilizing an online Nyquist calculator (https://svi.nl/NyquistCalculator) allowing for sub-diffraction limited resolution following image restoration. All images were rendered on a high performance CUDA-GPU enabled workstation. 3D renders were generated for volume-object analysis with Huygens Object Analyzer software, which was also used to quantitate overlapping volumes for identified objects.

### Proximity ligation assay

Proximity ligation assay (PLA) was performed according to manufacturer instructions (Millipore). PLA reports direct *in situ* interactions between proteins that are within 40nm of each other and revealed as fluorescent dots.

### High content imaging

All high content experiments were performed on Huh7 cells in 96-well plate format using optical-quality glass-bottomed plates. Imaging and analysis were performed using a Cellomics CellInsight CX7 scanner and driven by iDEV software (Thermo Fisher Scientific). Primary objects (cells, identified based on nuclear staining with Hoechst 33342 and regions of interest (ROI) were algorithm-defined by shape/segmentation, maximum/minimum average intensity, total area and total intensity to automatically identify puncta within valid primary objects. Five wells (>1000 cells/well) were analyzed per treatment per experiment. All data acquisition and analysis were computer driven and independent of human operators.

### Mitochondrial isolation experiments

Subcellular fractionation was performed with a QProteome mitochondria isolation kit (Qiagen) according to the instruction manual. In brief, 10^7^ Huh7 and HeLa cells were re-suspended in 1 mL of lysis buffer, incubated for 10 min at 4⁰ C and centrifuged at 1000x g for 10 min. The supernatant was transferred into a separate tube as cytosolic fraction, while the pellet was re-suspended in 1.5 mL of ice-cold disruption buffer, rapidly passed through 26g needle 10-15 times to disrupt cells and centrifuged at 1000x g for 10 min, 4⁰ C. The supernatant was then re-centrifuged at 6000x g for 10 min, 4⁰ C. The pellet obtained after centrifugation comprised the mitochondrial fraction.

### Luciferase assays

20000 HEK293T cells were plated in 96 well plates prior to transfection with the *Renilla* luciferase internal control reporter plasmid pRL-TK (thymidine kinase promoter dependent *Renilla* luciferase), plasmids encoding firefly luciferase responsive to NF-κB or ISRE and GFP, GFP-TRIM5α with or without myc-TBK1 expression plasmids as previously described (Saha et al., 2020). 40-48h after transfection, the plate was assayed using the Dual-Glo Luciferase Assay System according to the manufacturer’s instructions (Promega, E2920) and read using a Microplate Luminometer (BioTek, SYNERGY HTX Multi-Mode reader). Firefly luciferase readings were normalized to Renilla luciferase readings in each well, and the data are represented as fold-change relative to GFP alone.

### Peptide array analysis

TRIM27 peptide arrays were synthesized on cellulose membranes using MultiPrep peptide synthesizer (INTAVIS Bioanalytical Instruments AG, Germany). Membranes were blocked with 5% non-fat dry milk in TBST and peptide interactions were tested with GST and GST-GABARAP by overlaying the membrane with 1 μg/ml of recombinant proteins and incubation for 2 h at room temperature. Bound proteins were visualized with HRP-conjugated anti-GST antibody.

### GST pull-down assays

GST pulldown assays were performed by incubating immobilized GST or GST-tagged proteins with ^35^S-labeled *in vitro* translated proteins. All GST-tagged proteins were expressed in *Escherichia coli* SoluBL21 (Genlantis). GST fusion proteins were purified on glutathione-Sepharose 4 Fast Flow beads (GE Healthcare 17-5132-01). ^35^S-labeled proteins were synthesized *in vitro* using the TnT T7 coupled reticulocyte lysate system (Promega). Translation reaction products from 0.25 μg of plasmid DNA were incubated with GST-labeled proteins on glutathione-Sepharose beads in NETN-E buffer (50 mM Tris, pH 8.0, 100 mM NaCl, 1 mM EDTA, 0.5% Nonidet P-40) supplemented with cOmplete Mini EDTA-free protease inhibitor cocktail tablets (1 tablet/10ml) (11836170001, Roche) for 1 h at 4°C. The beads were washed five times with 400 μl of NETN-E buffer, boiled with 2× SDS-PAGE gel loading buffer with 1mM DTT, and subjected to SDS-PAGE. Gels were stained with Coomassie Brilliant Blue and vacuum-dried. ^35^S-labeled proteins were detected using a Fujifilm bioimaging analyzer BAS-5000 (Fuji) and quantifications were performed using Image Gauge software (Fuji).

### Statistical analysis

Data are expressed as means ± SEM (n>3). Where not otherwise indicated, data were analyzed with unpaired two-tailed t-tests or ANOVA with Tukey’s post hoc analysis. Analysis was performed using GraphPad Prism10. Statistical significance is defined as *, P < 0.05; **, P < 0.01; ***, P < 0.001; ****, P < 0.0001.

## Abbreviations

IP: immunoprecipitation
h: hour
KDa: kilodalton
CCCP: carbonyl cyanide 3-chlorophenylhydrazone
IVM: ivermectin
min: minute
WT: wild-type
KO: knock out
T5: TRIM5
DMSO: dimethyl sulfoxide
OPTN: optineurin
5Z: (5Z)-7-Oxozeaenol
ng: nanogram
Ub: ubiquitin
pp242: 2-(4-amino-1-isopropyl-1H-pyrazolo[3,4-d]pyrimidin-3-yl)-1H-indol-5-ol
LLOME: Leu-Leu methyl ester hydrobromide
siRNA: small interfering RNA
RING: really interesting new gene
CC: coiled-coil
GST: glutathione-S-transferase
MW: molecular weight
CBB: coomassie brilliant blue
mATG: mammalian autophagy-related protein.

## SUPPLEMENTARY FIGURE LEGENDS

**Supplementary Figure S1 related to Figure 1. TRIM5α recruits active pTBK1 to mitochondria following uncoupling with CCCP. (A)** HeLa cells stably expressing TRIM5α-HA were transfected with mCherry-Parkin and treated with CCCP for 90 minutes prior to fixation, labeling, and confocal microscopy. Arrows indicate regions where TRIM5α and pTBK1 colocalize with each other and are colocalized with or contiguous to Parkin signal. Three-dimensional reconstructions of the regions bounded by the dashed lines are shown to the right. **(B)** Confocal analysis of the overlap between pTBK1 and TIM23 in WT and TRIM5 KO Huh7 cells following 2 hour treatment with CCCP. The total volume of pTBK1 or TIM23 structures, as well as the overlapping volume, was determined by image analysis software. The graphs show the volume of overlapping voxels normalized to the total volume of either pTBK1 or TIM23. N = 3 independent experiments; > 5000 structures analyzed per marker per experiment. Data: mean + SEM; *P* values determined by Student’s t test; *, *P* < 0.05.

**Supplementary Figure S2 related to Figure 2. TRIM5α overexpression increases TBK1 abundance and activation. (A)** The effect of TRIM5α expression on the mitochondrial recruitment of the indicated autophagy proteins following CCCP treatment. WT or TRIM5 knockout Huh7 cells were transfected with GFP alone or GFP-TRIM5α and were treated or not with CCCP for 4 hours prior to harvest and purification of mitochondrial fractions. The numbers indicate the relative abundance of the indicated protein, normalized to TIM23. **(B)** Lysates from parental TRIM5 knockout or TRIM5-TBK1 double knockout Huh7 cells were subjected to immunoblotting as indicated. **(C, D)** Immunoblot analysis of the effect of TAK1 inhibitor 5z-7-oxozeanol (5Z) on CCCP-stimulated degradation of COXII in Huh7 cells (C). The relative abundance of COXII from three independent experiments was plotted in (D). **(E-H)** Lysates from HEK293T cells transfected with myc-TBK1 and either GFP-TRIM5α or GFP alone and treated or not with CCCP were subjected to immunoblotting with the indicated antibodies. The plots show the results from three independent experiments. **(I)** HEK293T cells were transfected with GFP or GFP-TRIM5α and plasmids encoding constitutively active *Renilla* luciferase and firefly luciferase under the control of an NF-κB responsive promoter. Samples were harvested 24 hours after transfection, and luciferase values determined. Plots show the relative signaling, which is determined by dividing a sample’s firefly luciferase signal by its *Renilla* luciferase signal. **(J)** Dual-luciferase based measurement of interferon signaling in HEK293T cells transfected with plasmids encoding GFP or GFP-TRIM5α, firefly luciferase under the control of an interferon-stimulated response element (ISRE) promoter, and the indicated amounts of myc-TBK1. *P* values determined by Student’s t test (panel I) or ANOVA with Tukey’s multiple comparison test; *, *P* < 0.05; **, *P* < 0.01; ***, *P* < 0.001; ****, *P* < 0.0001; †, not significant.

**Supplementary Figure S3 related to Figure 3. Impact of TRIM5α on TBK1 post-translational modifications. (A)** Coimmunoprecipitation analysis of TBK1 ubiquitination in WT and TRIM5 knockout HEK293T cells transfected with HA-ubiquitin and myc-TBK1 and treated with IVM for the indicated time points. **(B)** Coimmunoprecipitation analysis of TBK1 ubiquitination in HEK293T cells transiently transfected with myc-TBK1 and HA-UB K63 and treated as indicated for 1 h. **(C)** Lysates from HEK293T cells transfected with myc-TBK1 and GFP alone or GFP-TRIM5α (WT and E2 binding patch mutants) and treated or not with CCCP for 2 hours were immunoblotted as shown. Data from three independent experiments showing the relative abundance of pTBK1 or total myc-TBK1, normalized to actin, are shown in (**D**, **E)**. CCCP-treated samples are shown in shaded triangles and DMSO-treated samples are shown in open triangles. *P* values determined by ANOVA with Tukey’s multiple comparison test; **, *P* < 0.01; ***, *P* < 0.001; ****, *P* < 0.0001.

**Supplementary Figure S4 related to Figure 5. Mapping the determinants of TBK1 interactions with TRIM5α. (A)** Schematic of the TRIM5α deletion and substitution mutants used. Mapping experiments revealed that TBK1 interacted with the N-terminal region of TRIM5α. **(B)** HEK293T cells were transfected with plasmids encoding myc-TBK1 and GFP alone or GFP-TRIM5α (WT and mutants) and treated with CCCP for 2 hours prior to lysis and immunoprecipitation with anti-GFP. Red asterisks indicate the full-length GFP-tagged protein. **(C)** GST pull-down analysis of direct interactions between radiolabeled TBK1 and GST-TRIM5α. GST-NAP1 was used as a positive control for TBK1 binding. **(D)** HEK293T cells were transfected with GFP-TRIM5α and treated with CCCP for the indicated time points prior to lysis and immune precipitation with anti-GFP and immunoblotting with the indicated antibodies. **(E, F)** Assessment of K63-linked ubiquitination of GFP-TRIM5α from transfected HEK293T cells treated for the indicated time points with either IVM (E) or CCCP (F). **(G)** Coimmunoprecipitation analysis of interactions between GFP-TRIM5α and FLAG-SQSTM1 in HEK293T cells treated with CCCP for the indicated time points. **(H)** Coimmunoprecipitation analysis of interactions between GFP-TRIM5α and myc-TBK1 in WT HeLa cells or in HeLa cells lacking SQSTM1, NBR1, NDP52, OPTN, and TAX1BP1 (pentaKO) treated or not with IVM for 2 h.

**Supplementary Figure S5 related to Figure 6. Analysis of the role of the LIR motif in TRIM5α’s interactions with autophagy machinery and TBK1. (A)** Coimmunoprecipitation analysis of interactions between WT or ΔLIR2 GFP-TRIM5α and endogenous FIP200 in transfected HEK293T cells. **(B)** Coimmunoprecipitation analysis of interactions between WT or ΔLIR2 GFP-TRIM5α and FLAG-ATG13 in transfected HEK293T cells. **(C)** Lysates from HEK293T cells transfected with myc-ULK1 and GFP-tagged WT or ΔLIR2 TRIM5α were subjected to immunoprecipitation with anti-myc and immunoblotted as indicated. **(D)** Coimmunoprecipitation analysis of interactions between WT or ΔLIR2 GFP-TRIM5α and myc-ULK1 in WT HeLa cells or HeLa cells lacking all six mammalian Atg8 genes (HexaKO). **(E)** Coimmunoprecipitation analysis of interactions between WT or ΔLIR2 GFP-TRIM5α and mycTBK1 in transfected HEK293T cells treated with CCCP for 2 h. Red * indicate the position of the GFP-TRIM5α bands. **(F)** Lysates from HEK293T cells transfected with myc-TBK1 and GFP alone or GFP-TRIM5α (WT and ΔLIR2) and treated or not with CCCP for 2 hours were immunoblotted as shown. Plots show data from three independent experiments quantitating the relative abundance of pTBK1 or total myc-TBK1, normalized to actin. CCCP-treated samples are shown in shaded triangles and DMSO-treated samples are shown in open triangles. *P* values determined by ANOVA with Tukey’s multiple comparison test; ***, *P* < 0.001; ****, *P* < 0.0001.

**Supplementary Figure S6 related to Figure 7. Evaluation of the LIR motif in TRIM27 and TRIM27’s interactions with mammalian Atg8 proteins and TRIM5α. (A)** Alignment of TRIM27 and TRIM5α amino acid sequence, showing the two LIR motifs in TRIM5α (LIR1 and LIR2) and the homology between TRIM5α LIR2 and TRIM27 LIR. Asterisks indicate amino acids mutated to alanine in TRIM27^ΔLIR^. **(B)** Binding of GST-GABARAP to a TRIM27 peptide array. The entire TRIM27 protein sequence is tiled on the array with each dot representing a 2 amino acid shift in sequence from the neighboring dots. Bars indicate the sequence of the peptide spots that showed highest binding to GST-GABARAP. The highlighted sequence shows the residues present in all of the overlapping peptides. **(C, D)** GST pull-down analysis of interactions between *in vitro* translated ^35^S-labeled Myc-TRIM27^WT^ or Myc-TRIM27^ΔLIR^ against GST-tagged mAtg8 proteins. Plot shows quantitation of 3 independent experiments. Data, mean ± SD. *P* values determined by student’s T-test; *, *P* < 0.05; **, *P* < 0.001. **(E)** HeLa cells stably expressing TRIM5α-HA were transfected with plasmids encoding GFP-TRIM27 and mCherry-Parkin and the colocalization between GFP and HA was determined by confocal microscopy. mCherry signal was diffusely cytosolic (as expected) and is not shown. **(F)** Literature-based string schematic summarizing TRIM5α’s interactions with autophagy/mitophagy proteins. Blue lines between nodes indicate data from previous studies, green lines indicate data shown in this manuscript. Red arrows indicate interactions with TRIM5α that are enriched in response to mitochondrial damage.

## Notes

### Competing Interest Statement

The authors have declared no competing interest.

