## Supplementary figures and images for "TBK1 is ubiquitinated by TRIM5α to assemble mitophagy machinery"

### Figure S1

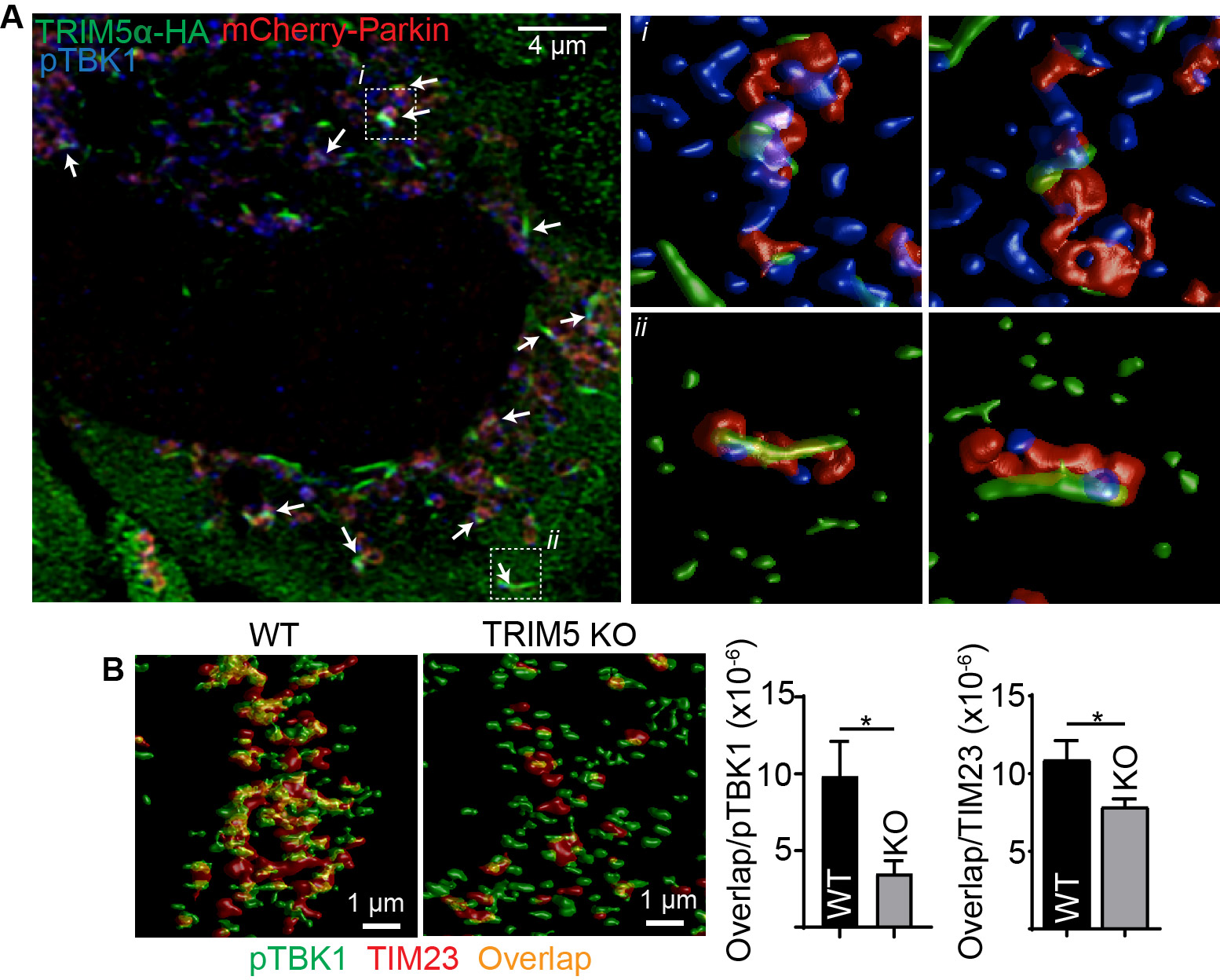

### Figure S2

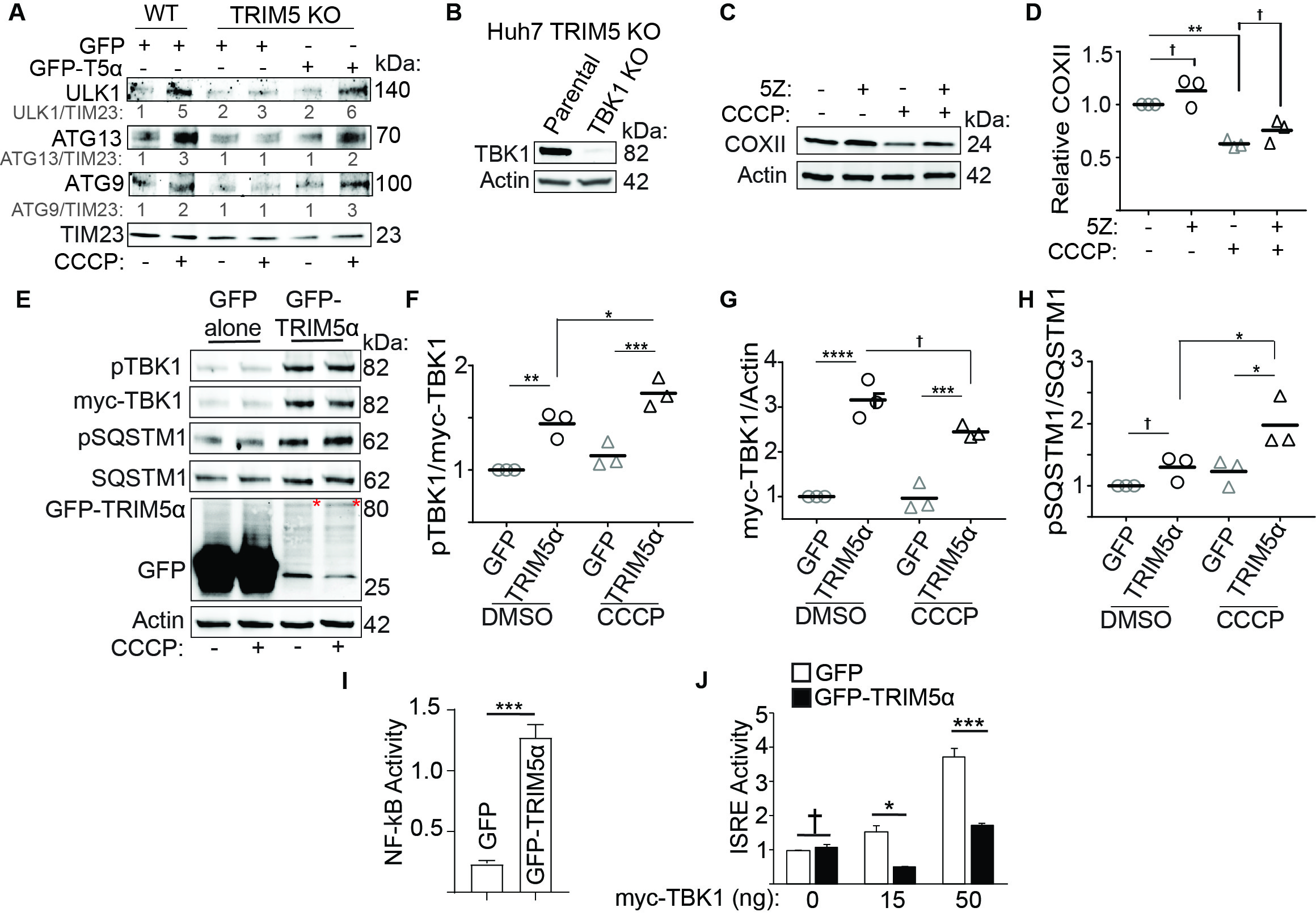

### Figure S3

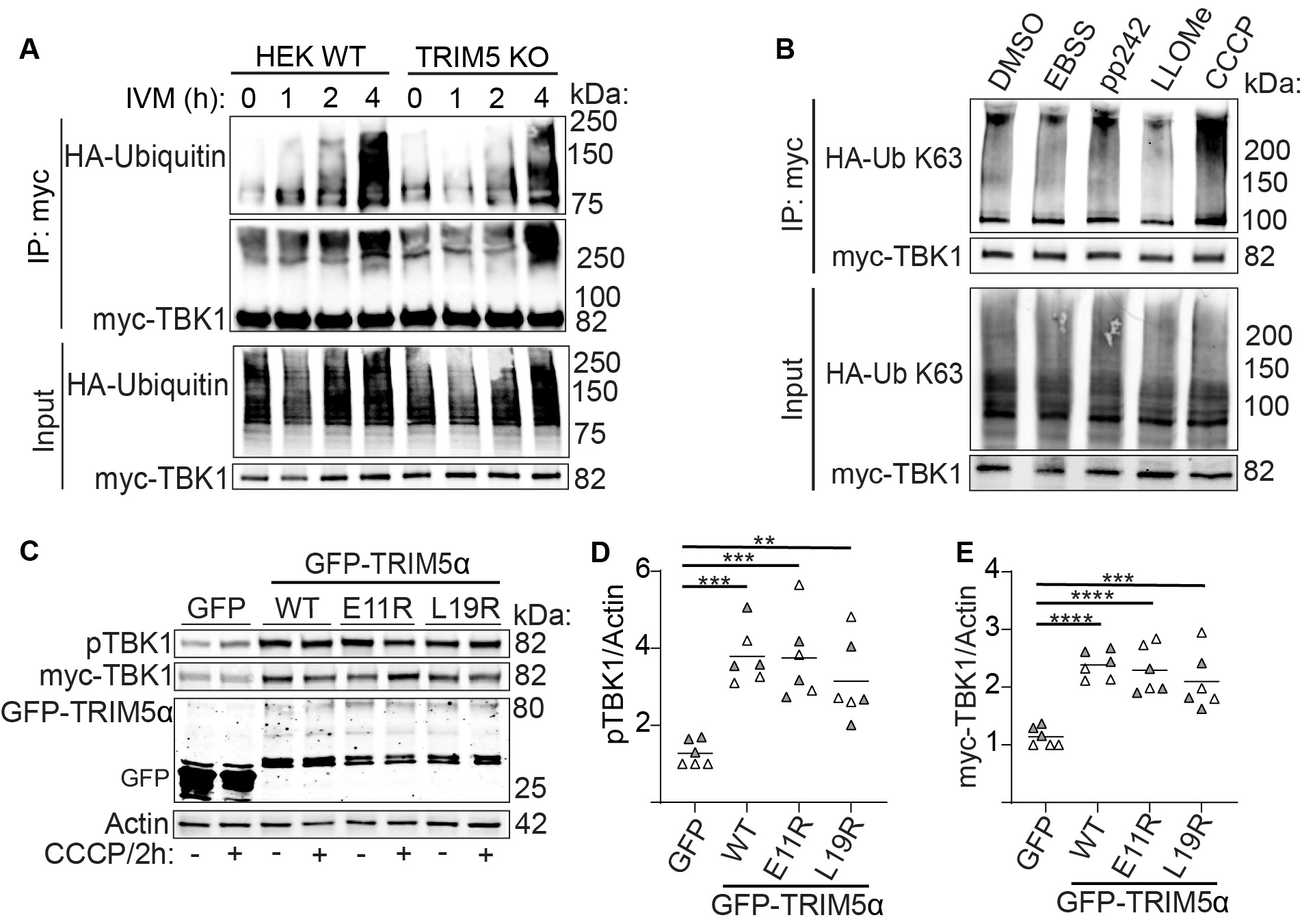

### Figure S4

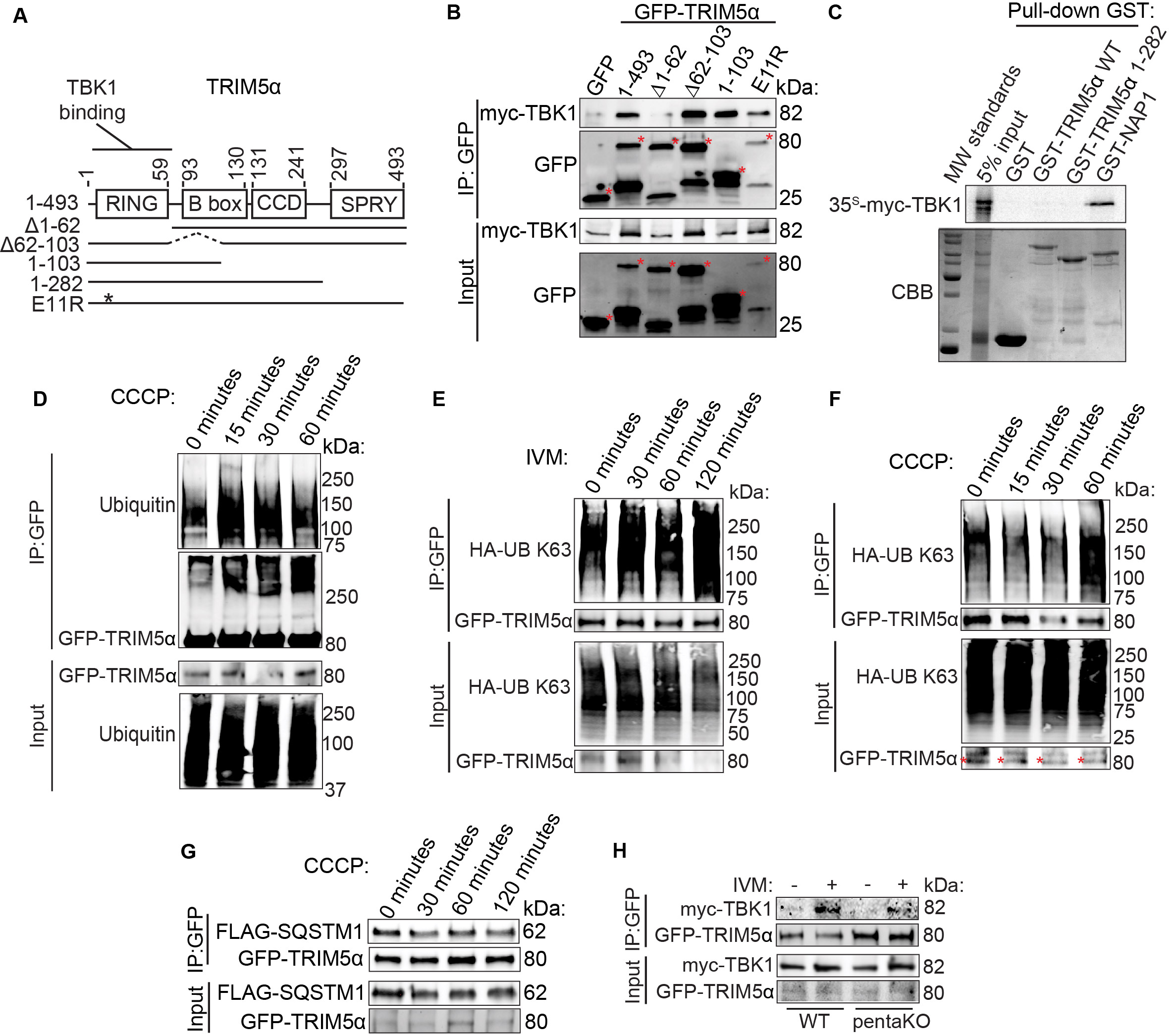

### Figure S5

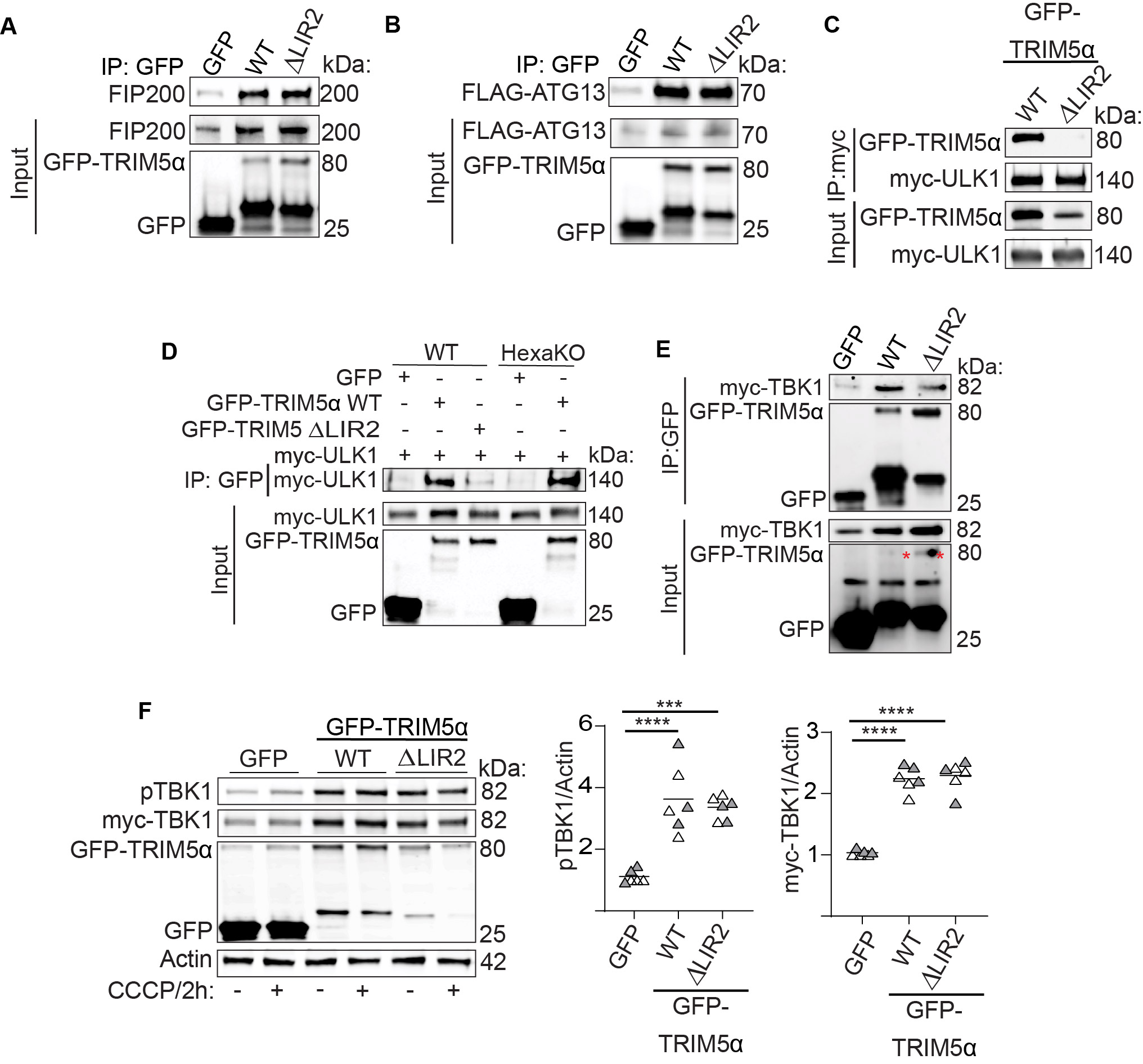

### Figure S6

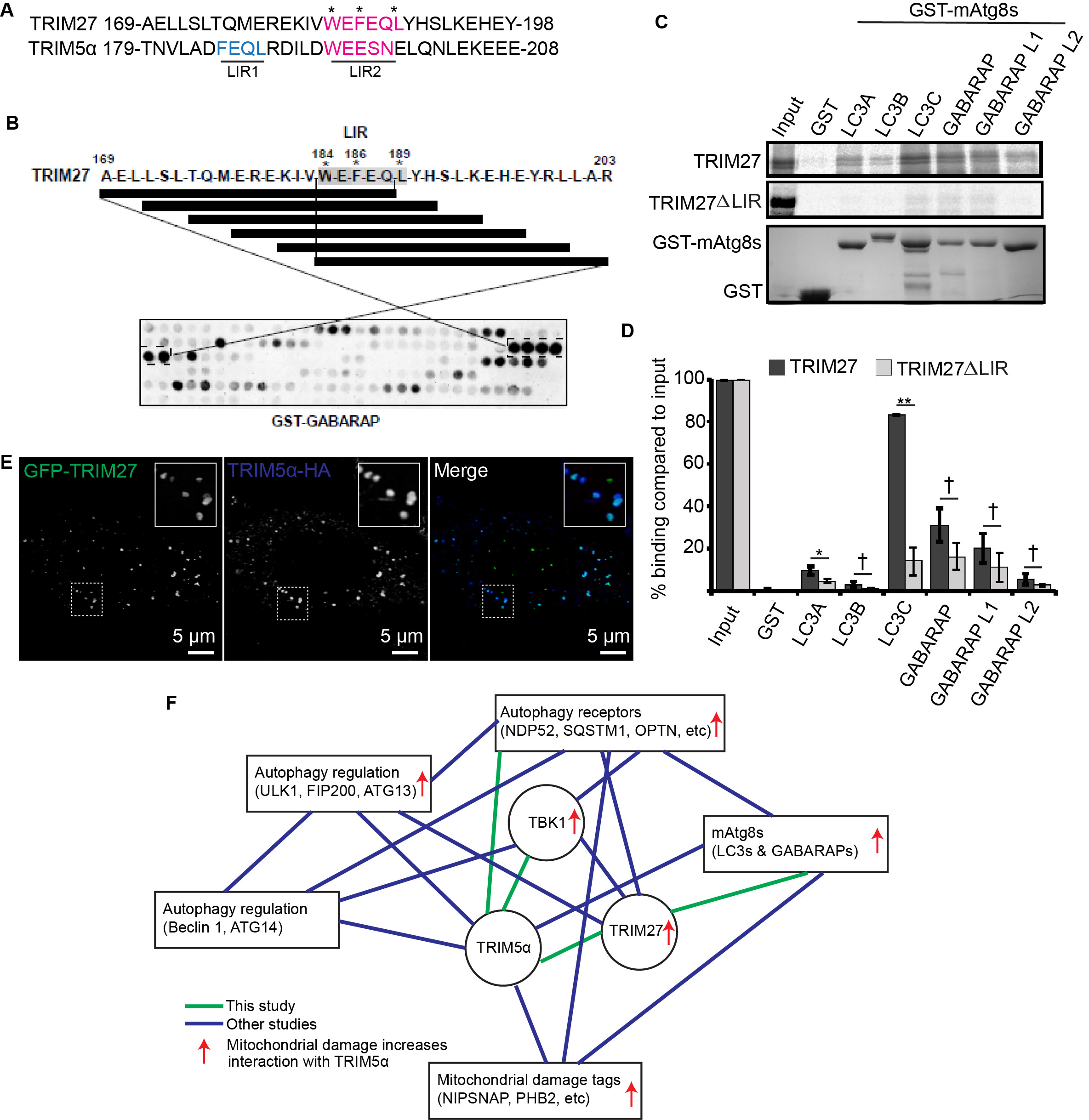
